# Phosphorylation Promotes DELLA Activity by Enhancing Its Binding to Histone H2A at Target Chromatin in *Arabidopsis*

**DOI:** 10.1101/2023.10.10.561786

**Authors:** Xu Huang, Rodolfo Zentella, Jeongmoo Park, Larry Reser, Dina L. Bai, Mark M. Ross, Jeffrey Shabanowitz, Donald F. Hunt, Tai-ping Sun

**Affiliations:** Department of Biology, Duke University, Durham, North Carolina 27708, USA; Department of Chemistry, University of Virginia, Charlottesville, Virginia 22904, USA; Department of Pathology, University of Virginia, Charlottesville, Virginia 22903, USA; U.S. Department of Agriculture, Agricultural Research Service, Plant Science Research Unit, Raleigh, NC 27607, USA; Department of Crop and Soil Sciences, North Carolina State University, Raleigh, NC 27695, USA; Syngenta, Research Triangle Park, NC 27709, USA

## Abstract

DELLA proteins are conserved master growth regulators that play a central role in controlling plant development in response to internal and environmental cues. DELLAs function as transcription regulators, which are recruited to target promoters by binding to transcription factors (TFs) and histone H2A via its GRAS domain. Recent studies showed that DELLA stability is regulated post-translationally via two mechanisms, phytohormone gibberellin-induced polyubiquitination for its rapid degradation, and Small Ubiquitin-like Modifier (SUMO)- conjugation to alter its accumulation. Moreover, DELLA activity is dynamically modulated by two distinct glycosylations: DELLA-TF interactions are enhanced by *O*-fucosylation, but inhibited by *O*-linked *N*-acetylglucosamine (*O*-GlcNAc) modification. However, the role of DELLA phosphorylation remains unclear. Here, we identified phosphorylation sites in REPRESSOR OF *ga1-3* (RGA, an AtDELLA) purified from *Arabidopsis* by tandem mass spectrometry analysis, and showed that phosphorylation of the RGA LKS-peptide in the poly- S/T region enhances RGA-H2A interaction and RGA association with target promoters. Interestingly, phosphorylation does not affect RGA-TF interactions. Our study has uncovered that phosphorylation is a new regulatory mechanism of DELLA activity.

## Introduction

The DELLA proteins (DELLAs) are nuclear-localized transcription regulators, which were initially identified as repressors of the phytohormone gibberellin (GA) signaling in *Arabidopsis thaliana* ^1,2^. Further studies show that DELLA orthologs are conserved master growth regulators in plants, which play a central role in coordinating multiple signaling activities in response to biotic and abiotic cues ^3–5^. DELLAs contain an N-terminal DELLA domain and a C-terminal GRAS domain ^1,2,6,7^. The DELLA domain functions as a regulatory sequence that interacts with the GA-bound receptor GIBBERELLIN INSENSITIVE1 (GID1), thereby promoting recruitment of the SCF^SLY1/GID2^ E3 ubiquitin ligase for polyubiquitination and proteolysis of the DELLA protein by the 26S proteasome ^8–15^. DELLAs mediate transcription reprogramming by direct interaction of their GRAS domain with hundreds of different transcription factors (TFs) ^3–5,16^. By characterizing new missense alleles of an *Arabidopsis DELLA*, *REPRESSOR OF ga1-3* (*RGA*), we recently revealed that formation of the TF-RGA-histone H2A complexes at the target chromatin is essential for RGA activity ^17^. The LHR1 subdomain within the RGA GRAS domain binds to TFs which then facilitate recruitment to target promoters by binding to TFs, while RGA- H2A interaction via the PFYRE subdomain, also within the GRAS domain stabilizes the TF- RGA-H2A complex at the target chromatin.

In addition to GA-dependent proteolysis mediated by polyubiquitination, DELLA activity is also modulated by several other post-translational modifications (PTMs) including Small Ubiquitin-Like Modifier (SUMO)-conjugation (SUMOylation), glycosylation, and phosphorylation ^18–21^. SUMOylated DELLA under salt-stress conditions was shown to sequester GID1 in a GA-independent manner, consequently increasing the amount of non-SUMO-DELLA and causing growth restriction ^22^. On the other hand, de-SUMOylation of DELLA under regular growth conditions promotes stamen filament elongation ^23^. Besides SUMOylation, recent genetic and biochemical studies revealed that DELLA activity is oppositely regulated by two distinct types of *O*-glycosylation of Ser and Thr residues; i.e., *O*-linked *N*-acetylglucosamine (*O*- GlcNAc) and *O*-fucose modifications ^19,20^. *O*-fucosylation of DELLA by SPINDLY (SPY) enhances DELLA binding to TFs (e.g., BZR1 and PIFs), whereas *O*-GlcNAcylation of DELLA by SECRET AGENT (SEC) reduces DELLA activity ^24,25^. It was proposed that *O*-Fuc and *O*- GlcNAc modifications may modulate DELLA activity and plant growth in response to nutrient availability as *O*-GlcNAcylation serves as a nutrient sensor in metazoans ^25,26^.

Besides SUMOylation and *O*-glycosylation, DELLA proteins are also phosphorylated. However, the precise role of phosphorylation in DELLA function is unclear as several studies have provided conflicting results. An earlier study showed that phosphorylation of the rice DELLA, SLENDER RICE1 (SLR1) does not affect F-box protein GID2 binding affinity, suggesting that GA-induced SLR1 degradation does not require its phosphorylation ^27^. However, another study suggests that phosphorylation of SLR1 by a casein kinase I (CK1), EARLIER FLOWERING1 (EL1), promotes its stability. The *el1* mutant flowers early and exhibits elevated GA response ^28^. Moreover, GA-induced degradation of SLR1-YFP in the *el1* mutant background was faster than in the WT background, suggesting that EL1 enhances DELLA stability. However, SLR1 phosphorylation by EL1 was only demonstrated in vitro, and phosphosites in SLR1 have not been identified. The potential role of phosphorylation has also been studied by mutating conserved Ser/Thr residues (to Ala or Asp/Glu) in *Arabidopsis* DELLAs, RGL2 or RGA and then monitoring the protein stability/activity in tobacco BY2 cells (RGL2) or in transgenic *Arabidopsis* (RGA) ^29,30^. Ala substitutions (RGA6A-GFP) appear to reduce RGA protein stability, whereas Asp mutations (RGA6D) stabilize RGA. Again, these results suggest that phosphorylation may enhance DELLA stability, although there is no evidence of phosphorylation of these S/T residues in planta. In addition, these Ser/Thr-to-Ala substitutions are located in the GRAS domain, and may affect RGA/RGL2 activity directly or affect SLY1 binding (hence stability).

To elucidate the role of phosphorylation in regulating DELLA function, it is crucial to identify DELLA phosphorylation sites in vivo and conduct functional analysis in planta. By affinity purification followed by MS/MS analysis, we have identified three phosphopeptides and mapped seven phosphosites in RGA using a transgenic *Arabidopsis* line carrying *P_RGA_:FLAG- RGA*. RGA phosphorylation was elevated under GA-deficient conditions. But GA treatment induced degradation of both phosphorylated and unphosphorylated RGA proteins, indicating that phosphorylation does not affect its stability. *rga^m2A^* encoding a mutant protein with eight Ser/Thr-to-Ala substitutions in the LKS-peptide (LKSCSSPDSMVTSTSTGTQIG) within the poly S/T region of RGA dramatically reduced its activity in planta. *rga^m2A^* did not confer changes in nuclear localization, GA-induced degradation or binding to RGA-interacting transcription factors [PHYTOCHROME INTERACTING FACTOR3 (PIF3) and BRASSINAZOLE-RESISTANT1 (BZR1)]. Importantly, co-IP and ChIP-qPCR assays showed that the rga^m2A^ mutant protein had decreased affinity to bind H2A, and reduced association with target promoters, revealing the mechanism of phosphorylation-induced RGA function.

## Results

### Elevated RGA phosphorylation under GA-deficient conditions

To investigate the role of DELLA phosphorylation, we first examined whether relative phosphorylation levels (vs. unphosphorylated form) of endogenous REPRESSOR OF *ga1-3* (RGA, an AtDELLA) are affected in different GA mutant backgrounds. We found that relative phospho-RGA (pRGA) levels were elevated under GA-deficient conditions (a GA-biosynthesis mutant *ga1-13* or WT Col-0 treated with GA biosynthesis inhibitor paclobutrazol [PAC]) compared to WT (**Fig. 1A**). Because phospho-RGA could not be separated clearly from the unphosphorylated form by standard SDS-PAGE, we used Phos-tag SDS-PAGE to detect phospho-RGA as the phosphate binding metal complex in the Phos-tag Acrylamide reagent causes retarded gel mobility of phosphorylated proteins ^31^. To monitor RGA phosphorylation more easily, we also analyzed FLAG-RGA in the *P_RGA_:FLAG-RGA ga1-13 della pentuple (ga1 dP)* transgenic line. Phosphatase treatment confirmed that the slower mobility band is phosphorylated FLAG-RGA (**Fig. 1B**). We further analyzed relative phospho-RGA levels in the F-box mutant *sly1-10*, which is a semi-dwarf with reduced GA responses because DELLAs accumulate to high levels ^11^. Due to the feedback mechanism that regulates GA biosynthesis, the *sly1* mutant contains elevated amounts of active GAs. The high GA content in *sly1* promotes inactivation of DELLAs by GID1 binding, which is proteolysis-independent ^32^. This is consistent with the less severe phenotype of the *sly1* mutant compared to *ga1*, although RGA accumulates to much higher level in *sly1* than in *ga1* (**Fig. 1C and 1D**). Phos-tag gel blot analysis showed that the relative phospho-RGA levels (pRGA vs. unphosphorylated form) in *sly1* were notably lower than those in *ga1* (**Fig. 1D**). These results indicate that RGA phosphorylation is elevated under GA deficiency (*ga1*) and reduced under high bioactive GA content (*sly1*). GA treatment induced degradation of both phosphorylated and unphosphorylated RGA proteins (**Fig. 1E**), indicating that phosphorylation does not alter RGA stability.

**Figure 1.**
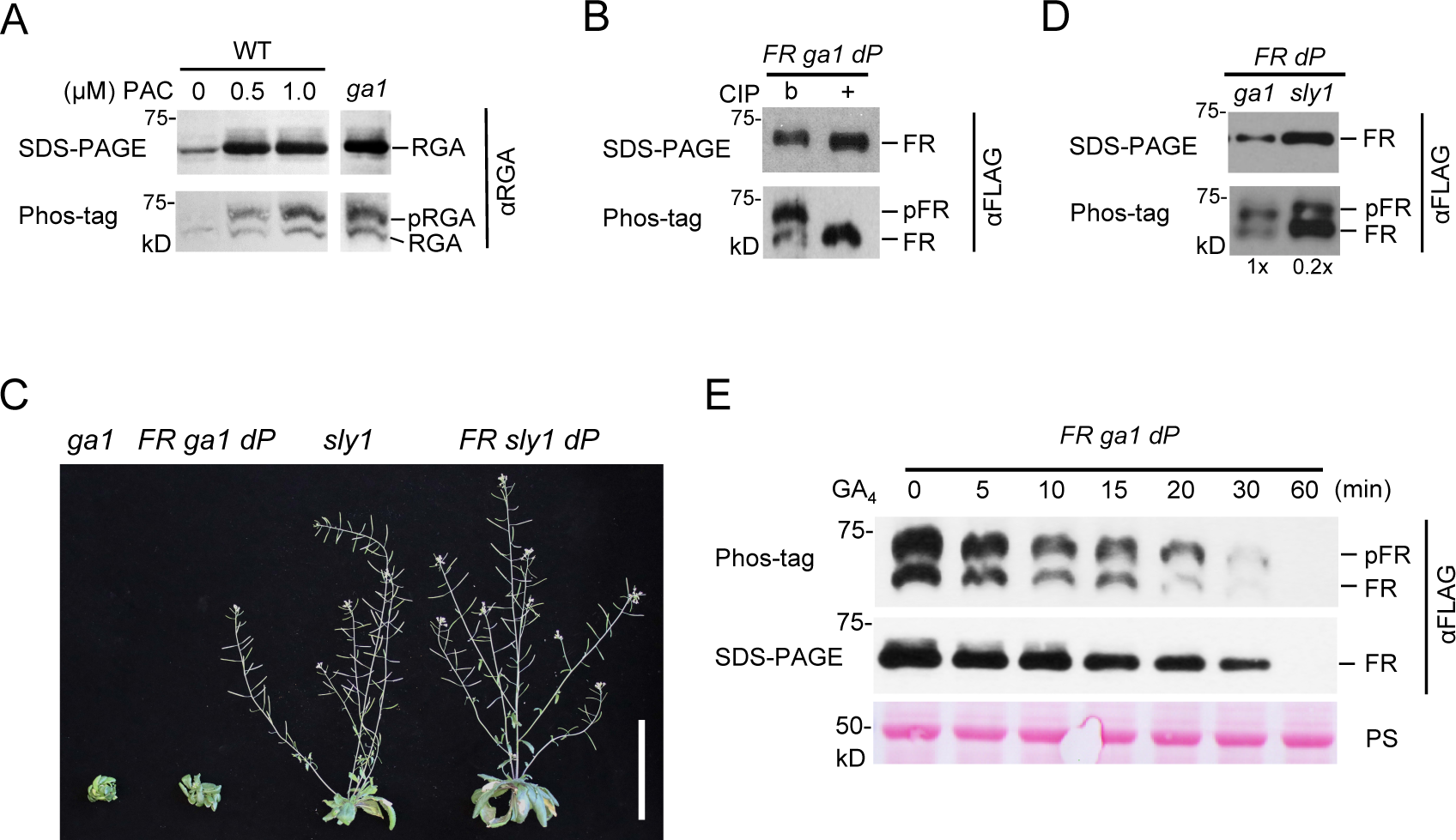
GA deficiency promotes RGA phosphorylation. **A)** Increased RGA phosphorylation by PAC treatment or *ga1* mutation. WT and *ga1* seedlings grown on media with different concentration of PAC for 10 days under long-day (LD) conditions. **B)** RGA phosphorylation pattern with or without CIP (Calf Intestinal Alkaline Phosphatase) treatment. b, boiled CIP. **C)** Representative 42d-old plants as labeled. *FR*, *P_RGA_:FLAG-RGA (FR)*. Bar = 3 cm. **D)** FLAG- RGA phosphorylation pattern in *FR ga1 dP* and *FR sly1 dP* lines. The blot contained total protein extracted from these lines. The *FR sly1 dP* protein sample was diluted 5-fold compared to that of *FR ga1 dP* because FLAG-RGA accumulated to very high levels in the *sly1* background. **E)** RGA phosphorylation did not affect GA-induced degradation. The protein blot contained total protein from *P_RGA_:FLAG-RGA ga1 dP* seedlings after 1 µM GA_4_ treatment for the indicated time. In A-B, D-E, proteins were analyzed by both standard SDS-PAGE and Phos- tag gels (containing 10 µM Phos-tag Acrylamide in A, B and E, and 25 µM in D), followed by immunoblotting with an anti-RGA antibody (in A) or anti-FLAG antibody (B, D and E). pRGA, phosphorylated RGA. pFR, phosphorylated FLAG-RGA. Ponceau S (PS)-stained blot in E indicated similar sample loading.

### Identification of RGA phosphorylation sites by LC-ESI-MS/MS

To identify phosphosites in RGA by MS analysis, we used transgenic *Arabidopsis* carrying *P_RGA_:FLAG-RGA^GKG^* in either *ga1-3 rga-24* or *sly1-10 rga-24* backgrounds. FLAG-RGA^GKG^ contains an extra trypsin cleavage site by inserting a Lys (K) residue within the Poly-S/T region that enables MS detection of this region ^24^ (**Fig. 2A**). FLAG-RGA^GKG^ is functional in planta to rescue the *rga* null phenotype. The affinity-purified FLAG-RGA^GKG^ samples from *ga1-3 rga-24* and *sly1-10 rga-24* backgrounds were analyzed by online liquid chromatography-electrospray ionization-tandem mass spectrometry (LC-ESI-MS/MS). Semi-quantitative analysis for relative peptide abundances was determined from ion currents detected in the MS1 survey scans (2-fold differences by this analysis may not be significant). Two highly phosphorylated RGA peptides, pep^LSN^ [LSNHGTSSSSSSISK(DK), 30.5%] and pep^LKS^ [(LK)SCSSPDSMVTSTSTGTQIGK], 28.6%), were identified in RGA^GKG^ purified from the *ga1* background (**Fig. 2A, 2B**, **Table 1**,

**Figure 2.**
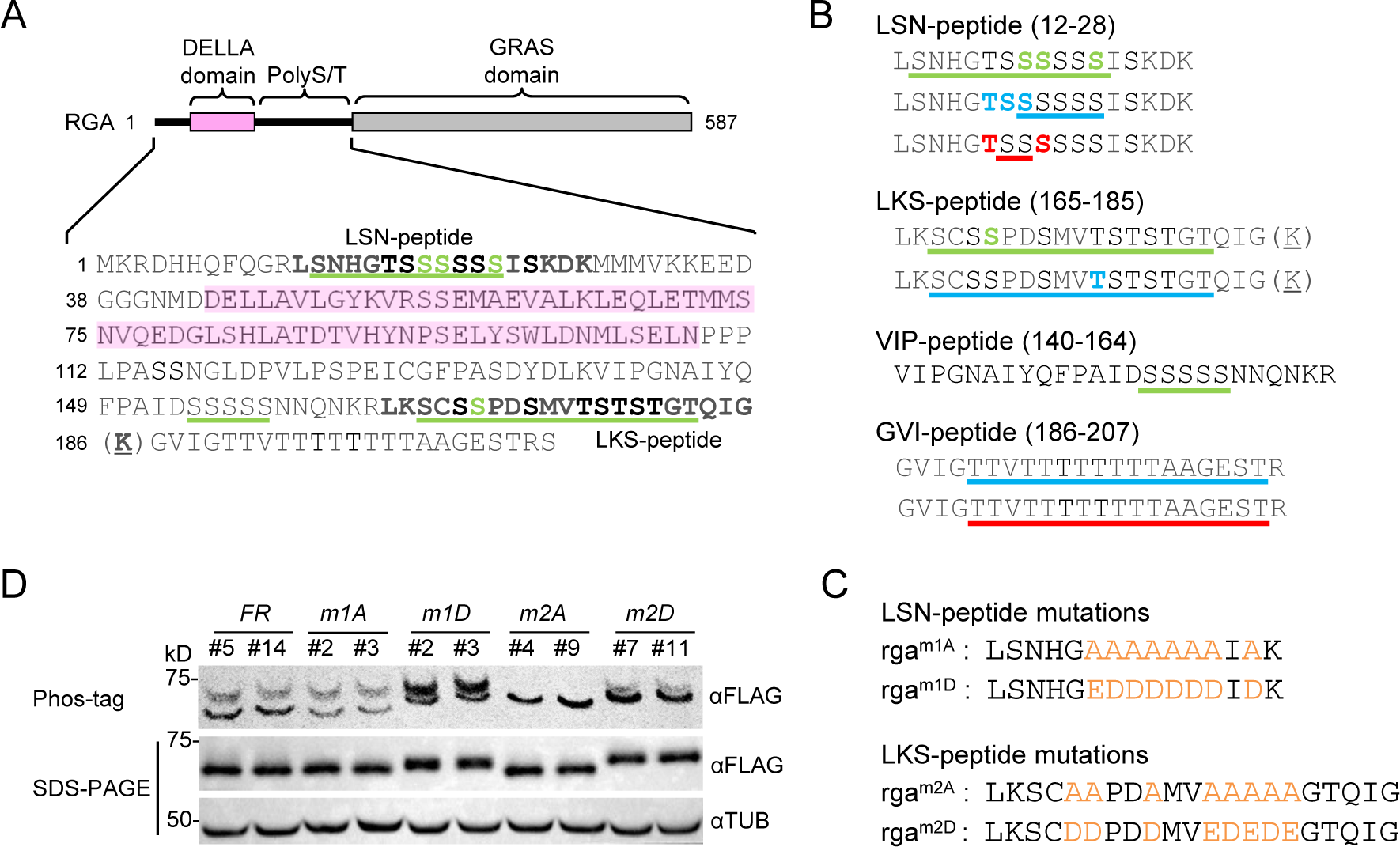
Identification of phosphorylation sites in RGA by LC-ESI-MS/MS. **A)** RGA phosphorylation sites identified by MS/MS. The schematic shows the RGA protein structure; two structurally disordered regions are indicated as solid black lines. The DELLA domain is shaded in pink. The pep^LSN^ and pep^LKS^ containing abundant phosphorylation are in bold (black letters). The S/T residues in green letters are confirmed phosphorylation sites. The underlined residues are amino acid stretches, in which one or more residues are modified (in addition to the identified sites), but the specific residues could not be mapped. The underlined K in parenthesis indicates the extra Lys residue in RGA^GKG^ for creating an additional trypsin cleavage site. **B)** Summary of PTM sites in RGA. Phosphorylation sites are labeled in green as described in A. *O*-fucosylation sites are labeled in red, *O*-GlcNAcylation sites in blue. The underlined K in parenthesis indicates the extra Lys residue in RGA^GKG^. **C)** Mutated residues in rga^m1A^, rga^m1D^, rga^m2A^ and rga^m2D^ are highlighted in orange. **D)** Expression and phosphorylation pattern of *P_RGA_:FLAG-RGA/-rga ga1 dP* transgenic lines using standard SDS-PAGE and Phos-tag (25 µM) gels. Protein blots were probed with anti-FLAG antibody or anti-tubulin (TUB, as a loading control).

**Table 1.**
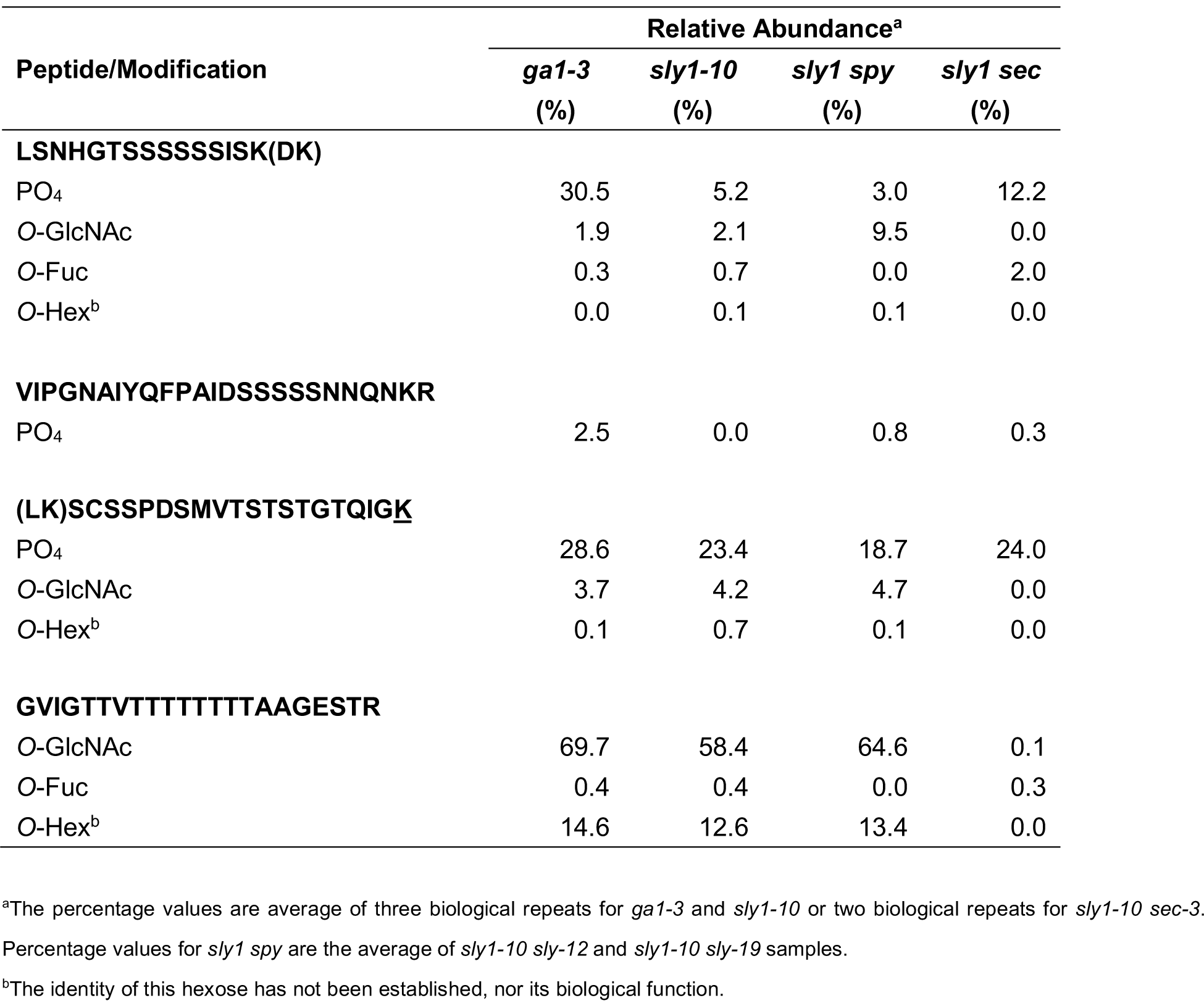
Relative Abundances of Posttranslational Modifications in FLAG-RGA^GKG^.

**Supplementary Table 1, and Supplementary Data 1-5**). pep^LSN^ is located in the poly-S region near the N-terminus, and pep^LKS^ is within the poly-S/T region downstream of the DELLA domain. Low levels of phosphorylation (2.5%) were also detected in another RGA peptide (pep^VIP^, VIPGNAIYQFPAIDSSSSSNNQNKR) that is immediate upstream of pep^LKS^. Interestingly, in the *sly1* background (with elevated GA), phosphorylation in pep^LSN^ was dramatically reduced (5.2% in *sly1* vs. 30.5% in *ga1*), while phosphorylation in pep^LKS^ was similar to that in *ga1* (23.4% in *sly1* vs. 28.6% in *ga1*, **Table 1**). MS analysis identified a total of seven phosphosites in RGA (**Fig. 2A, 2B** and **Supplementary Table 1**).

### RGA phosphorylation was not significantly altered by *spy* or *sec* mutation

The phosphorylation sites in RGA are located in the two disordered S- and S/T-rich sequences (flanking the DELLA domain, **Fig. 2A**), which were also shown previously to contain *O*-GlcNAc and *O*-Fuc sites, although most of the MS analyses were done using transiently expressed RGA in *N. benthamiana* ^24,25^. To investigate the interplay among these three PTMs in *Arabidopsis*, we identified RGA glycosylation sites and compared the relative abundances of each PTM in RGA^GKG^ purified from *ga-3*, *sly1-10*, *sly1-10 spy* (two *spy* alleles, *spy-12* and *spy- 19*, were included), and *sly1-10 sec-3* mutants. A summary of all PTM sites and their relative abundances is shown in **Fig. 2B, Table 1** and **Supplementary Table 1**. All three PTMs were detected in pep^LSN^, although *O*-GlcNAcylation and *O*-fucosylation were present at lower levels than phosphorylation. *O*-GlcNAcylation, but not *O*-fucosylation, was also identified in 4.2% of pep^LKS^. Additionally, we found pep^GVI^ (GVIGTTVTTTTTTTTAAGESTR) that contains the poly-T track was highly *O*-GlcNAcylated (69.7% in *ga1* and 58.4% in *sly1*), indicating that the levels of GlcNAcylation were not significantly altered by GA status. This is consistent with previous studies showing that transcript and protein levels of SPY and SEC are not regulated by GA ^24,25^. We detected only very low levels of *O*-fucosylation in pep^LSN^ and pep^GVI^. This is likely due to loss of *O*-Fuc during purification because no effective fucosidase inhibitors are available (in contrast to well characterized inhibitors for phosphatases and *O*-GlcNAcase). In summary, phosphorylation is located mainly in pep^LSN^ and pep^LKS^, but also at low levels in pep^VIP^. *O*- GlcNAcylation is highest in pep^GVI^ (poly-T track), but is also present at low levels in pep^LSN^ and pep^LKS^. *O*-Fuc was only detected at very low levels in pep^LSN^ and pep^GVI^ (**Fig. 2B**, **Table 1 and Supplementary Table 1**).

In pep^LSN^ that contains all three PTMs, phosphorylation was not affected by *spy*, although it was increased about 2-fold by *sec* (5.2% in *sly1* vs. 12.2% in *sly1 sec*) (**Table 1**). In contrast, phosphorylation levels in pep^LKS^ remained similar in the presence or absence of *spy* or *sec* mutation (**Table 1**), suggesting that phosphorylation of pep^LKS^ is not significantly affected by *O*- fucosylation or *O*-GlcNAcylation. *O*-GlcNAcylation in pep^LSN^ was elevated ∼5-fold by *spy* (2.1% in *sly1* vs 9.5% in *sly1 spy*), whereas *O*-fucosylation was increased ∼3-fold by *sec*, indicating antagonistic interaction between *O*-GlcNAc and *O*-Fuc modifications.

### Phosphorylation of pep^LKS^ enhanced RGA activity

To investigate the role of phosphorylation on RGA function, we generated transgenic *Arabidopsis* expressing four mutated rga proteins: rga^m1A^ (8 S/T-to-A in pep^LSN^), rga^m1D^ (8 S/T-to-D/E in pep^LSN^), rga^m2A^ (8 S/T-to-A in pep^LKS^), or rga^m2D^ (8 S/T-to-D/E in pep^LKS^) in the *ga1 dP* background (**Fig. 2C**). Three independent homozygous transgenic lines for each construct (*P_RGA_:FLAG-rga^m1A^*, *P_RGA_:FLAG-rga^m1D^*, *P_RGA_:FLAG-rga^m2A^*, *P_RGA_:FLAG-rga^m2D^*) with similar expression levels as the *P_RGA_:FLAG-RGA* lines were used for phenotype analysis (**Supplementary Fig. 1A-1B**). As expected, *FLAG-RGA* restored the dwarf phenotype in *ga1 dP*. Expression of *FLAG-rga^m1A^* only slightly increased growth to 14% of *ga1 dP*, indicating that *m1A* did not dramatically reduce RGA activity (**Fig. 3A-3B**, **Supplementary Fig. 1C-1D**). Phos-tag gel blot analysis showed that *rga^m1A^* did not reduce its phosphorylation (**Fig. 2D**). The phospho-mimic *FLAG-rga^m1D^*also slightly increased growth to 25% of *ga1 dP*, suggesting that *m1A* and *m1D* in pep^LSN^ may affect RGA activity independent of their phosphorylation status (**Fig. 3A-3B and Supplementary Fig. 1C-1D**). In contrast, *rga^m2A^* completely abolished phosphorylation of RGA, as detected by Phos-tag gel blot analysis (Fig. 2D), and markedly increased growth to 61% of *ga1 dP* (**Fig. 3A-3B and Supplementary Fig. 1C-1D**). The *rga^m2D^ ga1 dP* displayed similar dwarf phenotype as that of *FLAG-RGA ga1 dP*. These results support that *rga^m2A^* reduced RGA activity significantly. Hypocotyl elongation assays using representative *FLAG-RGA/rga* lines in the *ga1 dP* background further showed that only *FLAG-rga^m2A^* conferred much elevated GA responses comparing to *FLAG-RGA* (**Fig. 3C-3D**). We also compared activities of rga^m2A^, rga^m2D^ and RGA in regulating transcript levels of four selected RGA target genes, including two RGA-activated genes (*SCL3* and *GID1B*) and two RGA-repressed genes (*IAA16* and *EXP8*) by RT-qPCR analysis. Consistent with the whole plant phenotype results, FLAG-rga^m2A^ showed reduced activity while FLAG-rga^m2D^ displayed similar activity as FLAG-RGA in upregulating *SCL3* and *GID1B* (**Fig. 3E**), and downregulating *IAA16* and *EXP8* (**Fig. 3F**). These results indicate that although both pep^LSN^ and pep^LKS^ are highly phosphorylated, only pep^LKS^ phosphorylation promotes RGA activity.

**Figure 3.**
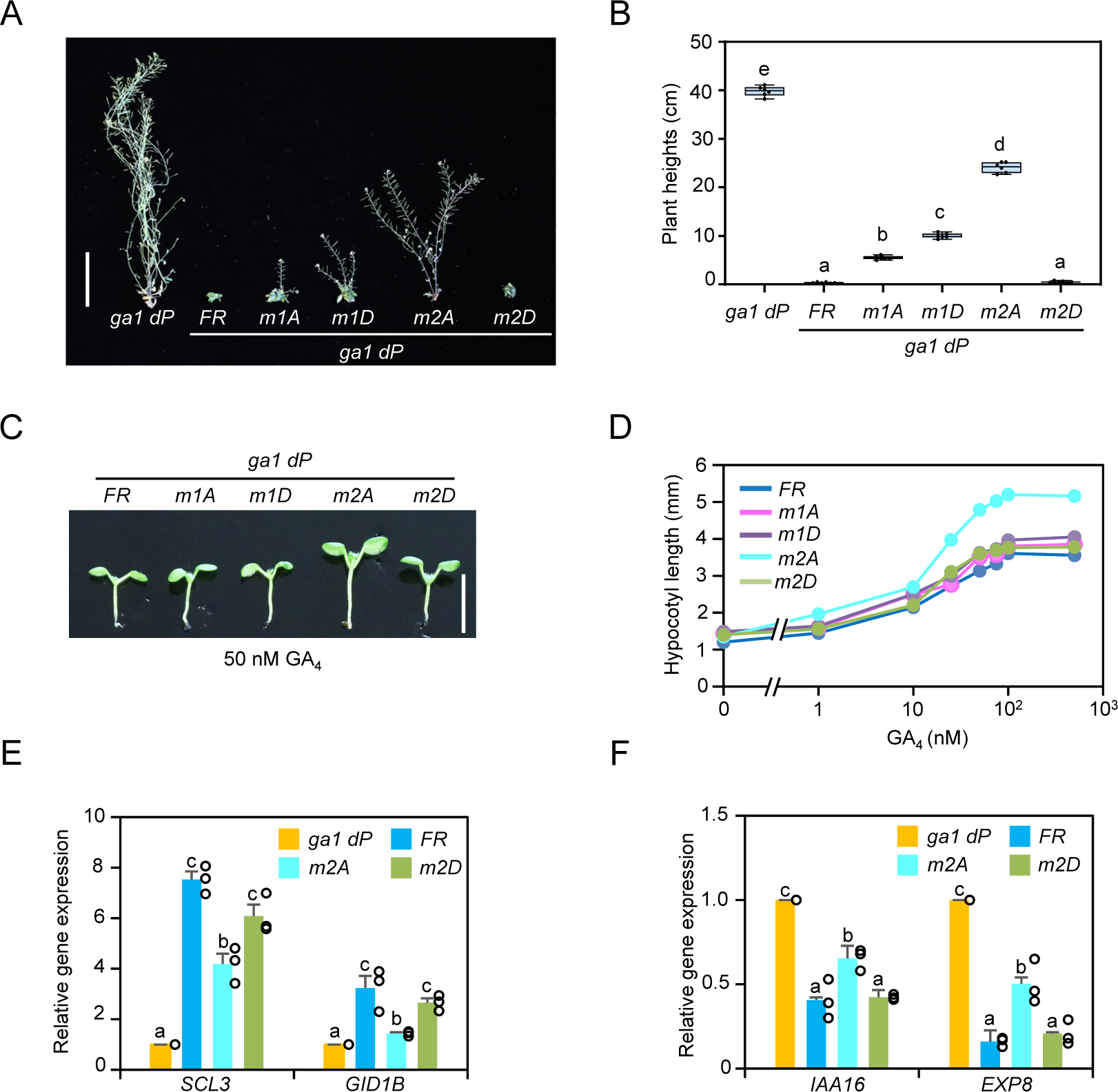
Phosphorylation of RGA in pep^LKS^ enhances RGA function**. A-B**) Phenotypes of FLAG-RGA/rga transgenic lines. In A, Representative 55-d-old plants under LD conditions. FR (#5), m1A (#3), m1D (#3), m2A (#4) and m2D (#7). Bar = 10 cm. In B, Boxplot showing final heights of different lines as labeled. n=6. Center lines and box edges are medians and the lower/upper quartiles, respectively. Whiskers extend to the lowest and highest data points within 1.5× interquartile range (IQR) below and above the lower and upper quartiles, respectively. Different letters above the bars represent significant differences (*p* < 0.01) as determined by two- tailed Student’s *t* tests. The phenotypic analysis was repeated three times with similar results. **C-D)** *FLAG-rga^m2A^ ga1 dP* displayed an enhanced GA response in hypocotyl growth. Seedlings were grown in medium containing 1 µM PAC and varying concentrations of GA_4_. Hypocotyl lengths were measured at day 9. Bar = 5 mm. In D, Average hypocotyl lengths are means ± SE. n =10-13. The assay was repeated three times with similar results. **E-F)** RT-qPCR showing rga^m2A^ caused reduced expression of RGA-induced genes (*SCL3* and *GID1B*) and increased expression of RGA-repressed genes (*IAA16* and *EXP8*). *PP2A* was used to normalize different samples. Means ± SE of three biological replicates are shown. Different letters above the bars represent significant differences (*p* < 0.05) by two-tailed Student’s *t*-test.

### Pep^LKS^ phosphorylation did not affect RGA protein stability or interaction with PIFs or BZR1

To investigate how pep^LKS^ phosphorylation enhances RGA function, we first examined the effects of rga^m2A^ and rga^m2D^ mutations on its subcellular localization and stability. Protein fractionation and immunoblot analysis showed that rga^m2A^ or rga^m2D^ did not alter nuclear localization or GA-induced degradation of RGA in *Arabidopsis* (**Supplementary Fig. 2A-2B**). Considering that *O*-Fuc and *O*-GlcNAc modifications oppositely alter RGA interactions with transcription factors PIFs and BZR1^24,25^, we tested whether rga^m2A^ reduced binding to PIF3 or BZR1. In vitro pulldown assays were performed using recombinant GST-tagged PIF3 and BZR1, and protein extracts from transgenic *Arabidopsis* expressing FLAG-RGA, FLAG-rga^m2A^ or FLAG-rga^m2D^ (**Fig. 4A**, **Supplementary Fig. 3A**). However, GST-PIF3 and GST-BZR1 pulled down FLAG-RGA, FLAG-rga^m2A^ and FLAG-rga^m2D^ similarly (**Fig. 4A**), suggesting that phosphorylation of pep^LKS^ regulates DELLA function differently from *O*-glycosylation.

**Figure 4.**
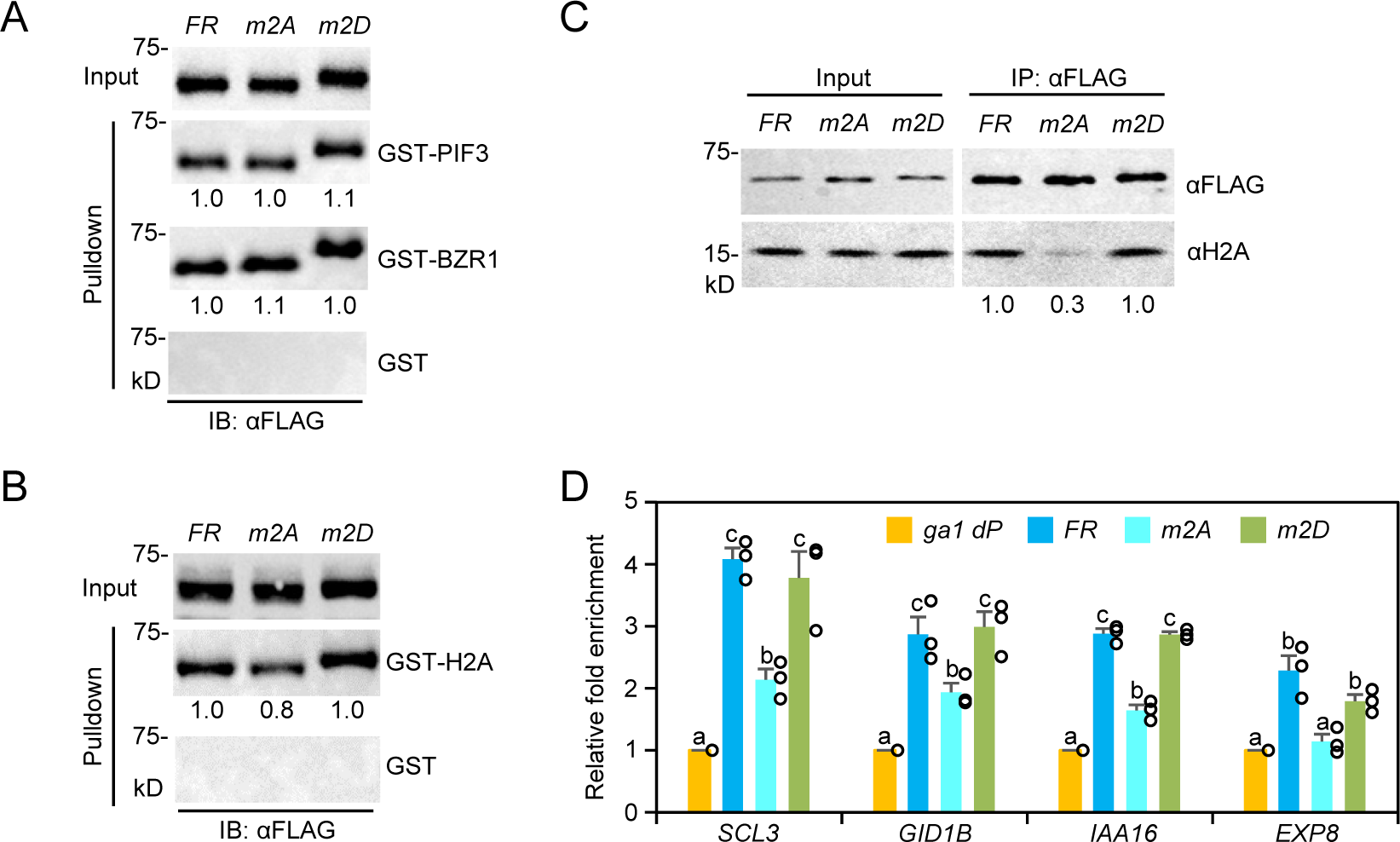
Phosphorylation promoted RGA interaction with H2A, but did not affect TF binding. **A)** In vitro pulldown assay showing rga^m2A^ and rga^m2D^ did not affect binding to PIF3 or BZR1. Recombinant GST, GST-PIF3 and GST-BZR1 bound to glutathione–Sepharose beads were used separately to pull down FLAG-RGA/rga from protein extracts from transgenic *Arabidopsis* in the *ga1 dP* background. Immunoblots containing input *Arabidopsis* extracts and pulldown samples were detected with an anti-FLAG antibody. PS-stained blots indicated that similar amounts of the GST/GST-fusion proteins were used in each set of the pulldown assays (**Supplementary Fig. 3A**). Representative images of two biological repeats are shown. **B-C**) rga^m2A^ reduced binding to H2A. In B, in vitro pulldown with GST or GST-H2A as described in (A). PS-stained blots indicated that similar amounts of the GST/GST-fusion proteins were used in each set of the pulldown assays (**Supplementary Fig. 3B**). In C, Co-IP assay showing the endogenous H2A was co-IP’ed more strongly by FLAG-RGA than by FLAG-rga^m2A^. FLAG-RGA/rga from protein extracts of transgenic *Arabidopsis* (in *ga1 dP* background) carrying *P_RGA_:FLAG-RGA/rga* were IP’ed using an anti-FLAG antibody. Immunoblots containing input *Arabidopsis* extracts and IP’ed samples were detected with anti-FLAG and anti-H2A antibodies, separately. Representative images of three (in B) or two (in C) biological repeats are shown. In A-B, relative amounts of FLAG-RGA/rga pulled down by GST fusion proteins are shown. The levels of FLAG-RGA were set as 1.0. In C, relative amounts of H2A co-IP’ed with FLAG-RGA/rga are shown. **D)** ChIP-qPCR analysis showing rga^m2A^ reduced association with target chromatin. ChIP was performed using transgenic lines containing *P_RGA_:FLAG-RGA*/*rga* in the *ga1 dP* background as labeled. The parental line *ga1 dP* was included as a control. Two RGA-activated genes (*SCL3* and *GID1B*) and two RGA-repressed genes (*IAA16* and *EXP8*) were tested by ChIP-qPCR using primers near the RGA binding peaks. The relative enrichment fold was calculated by normalizing against ChIP-qPCR of non-transgenic *ga1 dP* control using *PP2A*. Means ± SE of three biological replicates are shown. Different letters above the bars represent significant differences (*p* < 0.05) by two-tailed Student’s *t*-test.

### Pep^LKS^ phosphorylation increased H2A binding at target chromatin

We recently found that interaction with histone H2A is essential for RGA activity by promoting the formation of the TF-RGA-H2A complex at the target chromatin ^17^. Importantly, rga^m2A^ showed reduced affinity to H2A by in vitro pulldown as well as co-IP assays (**Fig. 4B, 4C**, and **Supplementary Fig. 3B**), supporting that phosphorylation of pep^LKS^ enhances RGA- H2A interaction in planta. Furthermore, ChIP-qPCR analysis was performed using transgenic lines containing *P_RGA_:FLAG-RGA* or *FLAG*-*rga* in the *ga1 dP* background, and showed that FLAG-rga^m2A^ significantly reduced association with four selected target promoters, including two RGA-activated genes (*SCL3* and *GID1B*) and two RGA-repressed genes (*IAA16* and *EXP8*) (**Fig. 4D**). In contrast, FLAG-rga^m2D^ displayed similar association with target promoters as FLAG-RGA. These results provide strong evidence for the promoting role of RGA pep^LKS^ phosphorylation in binding H2A at target chromatin.

## Discussion

In this study, we showed that phosphorylation of the RGA pep^LKS^ (in the poly-S/T region) enhanced RGA-H2A interaction and RGA association with target promoters, while it did not affect RGA interaction with transcription factors PIF3 and BZR1. This conclusion is based on (1) direct detection of RGA phosphorylation sites in planta by MS/MS analysis; (2) in vivo functional analysis of FLAG-RGA vs FLAG-rga mutant proteins; and (3) in vivo co-IP and ChIP-qPCR assays. We recently showed that the PFYRE subdomain within the RGA GRAS domain is essential for H2A binding, while the LHR1 subdomain is required for interaction with TFs ^17^. Considering that pep^LKS^ is located in the poly-S/T region linking the DELLA domain and the GRAS domain of RGA, we propose that phosphorylation of pep^LKS^ may induce a conformational change to make the PFYRE subdomain more accessible for H2A binding (**Fig. 5**). The rga^m2A^ containing eight S/T-to-A substitutions within pep^LKS^ mimics unphosphorylated RGA, whose conformation might interfere with PFYRE subdomain-H2A interaction (**Fig. 5**).

**Figure 5.**
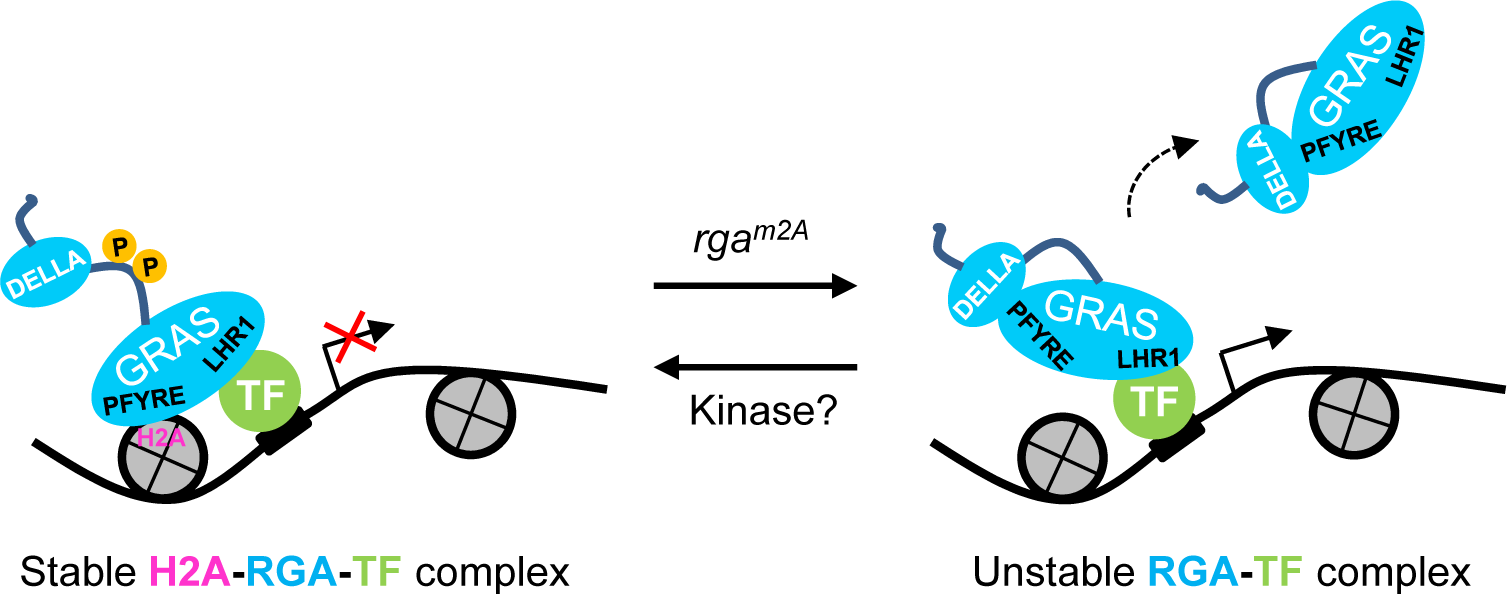
Proposed model for the regulatory role of RGA phosphorylation. RGA is recruited to target chromatin by interaction with TFs via the LHR1 subdomain, which is then stabilized by binding to H2A via its PFYRE subdomain to form H2A-RGA-TF complexes. Phosphorylation of the RGA pep^LKS^ within the poly S/T region (between the DELLA domain and the GRAS domain) by unidentified kinase(s) causes a conformational change that promotes RGA-H2A binding. The rga^m2A^ mutant protein abolishes RGA phosphorylation, and adopts a different protein conformation that interferes with H2A binding.

Our MS analysis showed that both pep^LSN^ and pep^LKS^ contained high levels of phosphorylation in the GA-deficient *ga1* background. However, phosphorylation of pep^LSN^ near the N terminus of RGA does not appear to play a significant role in regulating its activity because both *rga^m1A^* and *rga^m1D^* only slightly reduced RGA activity in planta. Intriguingly, pep^LSN^ showed much reduced phosphorylation in *sly1* (with elevated GA) compared to *ga1*. The reduced pep^LSN^ phosphorylation in *sly1* might be caused by increased GA-GID1 binding to the nearby DELLA domain in this mutant background.

The specific protein kinase(s) that phosphorylates *Arabidopsis* DELLA proteins remain elusive. A CK1 encoded by *EL1* in rice was reported to phosphorylate SLR1 (the rice DELLA protein), although evidence for this was only demonstrated in vitro ^28^. There are four *Arabidopsis* EL1-LIKEs (AEL1-4) ^33^, which are also known as MUT9-like kinases (MLKs) ^34^ and photo- regulatory protein kinases (PPKs) ^35^. To determine whether MLKs phosphorylate RGA in *Arabidopsis*, we introduced *P_RGA_:FLAG-RGA* into double and triple *mlk* mutants. However, our Phos-tag gel analysis did not detect any notable reduction in RGA phosphorylation in these mutants, suggesting that MLK1-4 do not play a major role in RGA phosphorylation (**Supplementary Fig. 4**). Consistent with our results, a recent phospho-proteomics study using *Arabidopsis AEL* overexpression lines and triple *ael* mutants did not identify any DELLA proteins as the substrates of these kinases ^36^. While preparing our manuscript, a GSK3/SHAGGY-like kinase-encoding gene *GSK3* in *Triticum aestivum* (wheat) was reported to phosphorylate DELLA (Rht-B1b) ^37^, although Rht-B1b phosphorylation by GSK3 has not been demonstrated in planta. Three phosphorylation sites in Rht-B1b located between the DELLA and GRAS domains were identified by in vitro enzyme reactions in the presence of GSK3, followed by MS analysis. Ser-to-Ala substitutions at all three phosphorylation sites led to reduced Rht- B1b activity in transgenic wheat, which is consistent with our finding that Ala substitutions in RGA pep^LKS^ reduced RGA activity. However, the in vitro protein degradation assay further suggested that phosphorylation also stabilizes Rht-B1b ^37^. This is in contrast to our results showing that Ala substitutions in RGA pep^LKS^ did not alter its stability in planta. GSK3 in wheat is an ortholog of BRASSINOSTEROID INSENSITIVE 2 (BIN2) in *Arabidopsis*^37^, which is a negative regulator of brassinosteroid (BR) signaling, and BR activates its signaling pathway by inducing BIN2 degradation^38^. We previously showed that BR treatment does not reduce RGA stability^39^ or phosphorylation levels in *Arabidopsis* (not shown), suggesting that RGA is unlikely to be phosphorylated by BIN2.

In summary, our study has uncovered a key role of phosphorylation in enhancing DELLA activity by promoting DELLA-H2A interaction at target chromatin. This adds a third mechanism to modulate DELLA activity, in addition to protein stability (by ubiquitination and SUMOylation) and binding affinity to TFs (by *O*-GlcNAc and *O*-Fuc modifications). Identification of protein kinase(s) for RGA pep^LKS^ phosphorylation will help reveal the internal and/or external cues that trigger DELLA phosphorylation. Furthermore, structural analysis of phosphorylated vs. unphosphorylated RGA will help to further elucidate the molecular mechanism of DELLA-mediated transcription reprogramming.

## Methods

### Plant materials, growth conditions, and generation of transgenic lines

In most experiments, *Arabidopsis* plants were grown in the growth room under long-day (LD) conditions (16 h light, 22 °C; 8 h dark, 20 °C). The *ga1-13*, *sly1-10* (bc 6x Col-0), and *ga1-13 della pentuple* (*ga1 dP*) are in the Col-0 background ^17,40^. The *ga1-3 rga-24* and *sly1-10 rga-24* double mutants are in the L*er* background ^24^. The *mlk1* (SALK_026482), *mlk2* (SALK_035080), *mlk3* (SALK_017102) and *mlk4* (SALK_1615), all in the Col-0 background, were obtained from Arabidopsis Biological Resource Center (https://abrc.osu.edu/). The double and triple homozygous *mlk* mutants were generated by crosses. Transgenic *Arabidopsis* lines, *P_RGA_:FLAG- RGA ga1-13 della pentuple (ga1 dP)*, *P_RGA_:FLAG-RGA^GKG^* in *ga1-3 rga-24*, *sly1-10 rga-24* or *sly1-10 sec-3 rga-24,* were generated previously ^17,24,25^. P*_RGA_:FLAG-RGA^GKG^* in *sly1-10 spy-12 rga-24 or sly1-10 spy-19 rga-24* were generated by crosses between *P_RGA_:FLAG-RGA^GKG^* in *sly1-10 rga-24* and different *spy* alleles. pRGA-His-3xFLAG-RGA, pRGA-His-3xFLAG-m1A, pRGA-His-3xFLAG-m1D, pRGA-His-3xFLAG-m2A, pRGA-His-3xFLAG-m2D constructs were introduced into *ga1-13 dP* by agrobacterium-mediated transformation. Independent transgenic lines with single insertion were selected by Basta resistance. Multiple independent transgenic lines (6 to 9) for each construct were screened by standard SDS-PAGE gel blot analysis to select for lines that expressed FLAG-RGA or -rga protein at similar levels. The *P_RGA_:FLAG-RGA* transgenic lines in the WT, *mlk* double and triple mutant backgrounds were generated by transformation. Transgenic lines for each genetic background that expressed similar levels of FLAG-RGA were used for further analysis.

### Plasmid construction

Primers and plasmid constructs are listed in **Supplementary Tables 2 and 3**, respectively. All DNA constructs generated from PCR amplification were sequenced to ensure that no mutations were introduced. Construction of pRGA-His-3xFLAG-RGA/m1A/m1D/m2A/m2D were generated using four constructs: J035 (RGA promoter, 8.1kb), pBm43GW ^41^, J015(RGA 3’UTR) and pDONR207-His-3XFLAG-RGA/m1A/m1D/m2A/m2D by Gateway LR reaction.

### Phenotype and statistical analyses

For final height measurement, the seeds of parental line *ga1 dP*, and transgenic lines carrying *P_RGA_:FLAG-RGA, P_RGA_:FLAG-rga^m1A^*, *P_RGA_:FLAG-rga^m1D^*, *P_RGA_:FLAG-rga^m2A^*, or *P_RGA_:FLAG-rga^m2D^* (all in the *ga1 dP* background) were treated with 10 µM GA_4_ for 3 days at 4 °C, washed 6 times with water, and then were sown in soil under LD. The experiment was repeated 3 times with similar results. For hypocotyl elongation analysis, surface-sterilized seeds were treated with 10 µM GA_4_ for 3 days at 4 °C, washed 6 times with water, and were plated on 0.5x Murashige and Skoog (MS) medium supplemented with different concentrations of GA_4_ for 9 days under LD conditions (16 µmol m-2 s-1 white light). Hypocotyl lengths were measured using ImageJ software (http://rsb.info.nih.gov/ij). Each experiment was performed at least three times with similar results and one set of representative results is shown. All statistical analyses were performed using Excel, and significant differences determined by Student’s t-tests.

### Reverse transcription (RT)-quantitative PCR (qPCR) and immunoblot analyses

Total RNA was extracted using the Quick-RNA MiniPrep kit (Zymo Research), and Reverse transcription was performed using Transcriptor First Strand cDNA Synthesis Kit (Roche) with anchored oligo dT_18_. RT-PCR analysis was performed using the FastStart Essential DNA Green Master mix and LightCycler 96 (Roche Applied Science). Relative transcript levels were determined by normalizing with *PP2A* (At1g13320) ^42^. Primers for the qPCR are listed in **Supplementary Table 2**. Primers for *SCL3*, *GID1B* and *EXP8* were reported previously ^40^.

For immunoblot assays, total proteins were extracted from 10 days–old seedlings using the 2×SDS extraction buffer (125 mM Tris-HCl pH 8.8, 4% SDS, 20% glycerol, 20% 2- Mercaptoethanol). Immunoblot analyses were performed using rat anti-RGA antiserum (DUR18, 1:2,000) ^11^, horseradish peroxidase (HRP)-conjugated anti-FLAG M2 mouse monoclonal (Sigma Aldrich A8592, 1:10,000 dilution), rabbit anti-histone 3 polyclonal antibody (Abcam ab1791, 1:5,000 dilution), mouse anti-tubulin antibody (Sigma T5168, 1:100,000) and rabbit anti-H2A polyclonal antibody (Millipore ABE327, 1:1,000 dilution). HRP-conjugated donkey anti-mouse IgG (Jackson ImmunoResearch #715-035-150) was used for anti-tubulin at a 1:10,000 dilution. HRP-conjugated goat anti-rabbit IgG (Thermo-Fisher #31462) was used to detect anti-H2A and anti-H3 at 1:10,000 dilution. HRP-conjugated goat anti-rat IgG (Pierce #31470) was used for anti-RGA (DUR18) at 1:10,000 dilution. Chemiluminescent signals were detected by iBright FL1500 (Invitrogen).

### Phos-tag mobility shift assay

Total proteins of 10-days-old seedlings were extracted from ground samples using the 2×SDS extraction buffer. The proteins were separated in a 6% SDS–PAGE gel containing 10-25 µM Phos-tag Acrylamide reagent (FUJIFILM Wako Chemicals USA Corp. #AAL-107) and 50 mM MnCl_2_. After electrophoresis, the gel was washed 3 times with transfer buffer without methanol plus 5 mM of EDTA, rinsed once with transfer buffer, and then transferred to nitrocellulose membranes for immunoblot analysis.

### In vitro pulldown and co-IP assays

In vitro pulldown assay was performed following the procedures published previously ^17,24^. Recombinant proteins (GST, GST-BZR1, GST-PIF3 and GST-H2A) expressed in BL21- CodonPlus (DE3)-RIL (Agilent Technologies) were purified using glutathione beads. GST and GST-fusion proteins bound to glutathione beads were then used separately to pull down FLAG-RGA/rga from protein extracts of transgenic *Arabidopsis* (in *ga1 dP* background) carrying *P_RGA_:FLAG-RGA/rga*.

For in vivo co-IP assays, total protein complexes were extracted from *Arabidopsis* and IP’ed using anti-FLAG-M2-Agarose beads as described ^17^. Samples were analyzed by SDS-PAGE and immunoblotting using anti-FLAG-HRP antibody (Sigma-Aldrich, 1:10,000) and anti- H2A antibody (Millipore; #ABE327). Quantitative analysis of the relative signal intensity was performed using ImageJ software (http://rsb.info.nih.gov/ij).

### Protein purification for MS analysis

His-FLAG-RGA was purified from P*_RGA_:FLAG-RGA^GKG^* transgenic *Arabidopsis* lines with *ga1- 3*, *sly1-10*, *sly1-10 sec-3*, *sly1-10 spy-12* or *sly1-10 spy-19 (*all in the *rga-24* background), following the tandem affinity purification procedures described previously^25^.

### Identification of PTM sites by online liquid chromatography tandem MS (MS/MS) analyses

Affinity-purified His-FLAG-RGA^GKG^ proteins extracted from *Arabidopsis* were trypsin-digested, and peptides were analyzed by online LC-electrospray ionization (ESI) tandem MS [electron-transfer dissociation (ETD) and collisionally activated dissociation (CAD) MS/MS] using a Thermo™ Orbitrap Fusion™ Tribrid™ mass spectrometer equipped with ETD ^25,43^. MS1 spectra were acquired in the Orbitrap with a resolution of 120,000, followed by low resolution data dependent MS2 analysis. Precursors of charge 2-6 were fragmented by CAD (30% normalized collision energy), and precursors of charge state 3-6 were fragmented using ETD with calibrated charge-dependent reaction times. Dynamic exclusion was included (repeat count of 1, repeat duration of 30 sec, exclusion duration of 10 sec).

Protein Metrics Byonic™ ^44^ was used to search data against a database containing the UniProt Reviewed ^45^ entries for *Arabidopsis thaliana* proteins with the addition of the sequence for 6His-3xFLAG-RGA^GKG^. Search settings included fully specific tryptic digestion, 3 potential missed cleavages, 10 ppm precursor mass tolerance, and 0.35 Da fragment mass tolerance. Alkylation of Cys residues was a fixed modification. Variable modifications included phosphorylation of Ser, Thr, and Tyr, *O*-fucosylation of Ser and Thr, *O*-GlcNAcylation of Ser and Thr, *O*-hexosylation of Ser and Thr, oxidation of Met, and the absence of alkylation on Cys. No manual cutoff based on false discovery rate or peptide score was applied. Byonic peptide- MS2 spectra matches were manually validated using both MS1 and MS2 spectra, and the modification site localization was confirmed by manual inspection of the MS2 spectra. Each peptide was quantified by integrating peak areas for all detected charge states including 13C isotopes for the two most abundant charge states.

### Chromatin immunoprecipitation (ChIP)-qPCR

Transgenic *Arabidopsis* seedlings carrying pRGA-His-3xFLAG-RGA, pRGA-His-3xFLAG- m2A and pRGA-His-3xFLAG-m2D (in the *ga1 dP* background) grown for 10 days were harvested and cross-linked in 1% formaldehyde solution for 20 min. ChIP-qPCR assay was performed using anti-FLAG-M2-Agarose beads (Sigma-Aldrich A2220) as described ^17^. The relative enrichment was calculated by normalizing against *ga1 dP* control samples using *PP2A* ^42^. Primers for the ChIP–qPCR are listed in **Supplementary Table 2**. Primers for *SCL3*, *GID1B*, *IAA16* and *EXP8* were reported previously ^17^.

### Accession numbers

Sequence information for *Arabidopsis* genes included in this article can be found in the GenBank/EMBL data libraries under accession numbers *RGA (*AT2G01570), SCL3 (AT1G50420), *GID1B (*AT3G63010), *IAA16* (AT3G04730), *EXP8 (*AT2G40610*). PP2A (*AT1G13320), *MLK1* (At5g18190), *MLK2* (At3g03940), *MLK3* (At2g25760), *MLK4* (At3g13670)

## Data Availability

The mass spectrometry proteomics data have been deposited to the ProteomeXchange Consortium via the PRIDE ^46^ partner repository with the dataset identifier XXXXX [Project DOI: XXXX].

## Acknowledgements

We thank Melissa Leyden for preparing MS figures and Emily Zahn and Ellen Speers for conducting the MS analyses. We also thank Neil Olszewski for providing plasmids JO15 and JO35. This work was supported by the National Institutes of Health (GM100051 to TPS, and GM037537 to DFH), and the National Science Foundation (MCB-1818161 to TPS).

## Author Contributions

T-pS, RZ and JP conceived and designed the research project. XH, RZ and JP performed molecular biology, genetics and biochemical analyses, and XH, JP, RZ and T-pS analyzed the data and generated figures. LR, DLB, MMR, JS and DFH analyzed the MS/MS data. T-pS and XH wrote the manuscript with input from all co-authors.

## ORCID

Xu Huang: orcid.org/0000-0002-2711-9920

Rodolfo Zentella: orcid.org/0000-0003-3986-6250

Jeffrey Shabanowitz: orcid.org/0000-0001-5750-3539

Donald F. Hunt: orcid.org/0000-0003-2815-6368

Tai-ping Sun: orcid.org/0000-0001-5223-2936

