## Supplementary Figures for "Phosphorylation Promotes DELLA Activity by Enhancing Its Binding to Histone H2A at Target Chromatin in *Arabidopsis*"

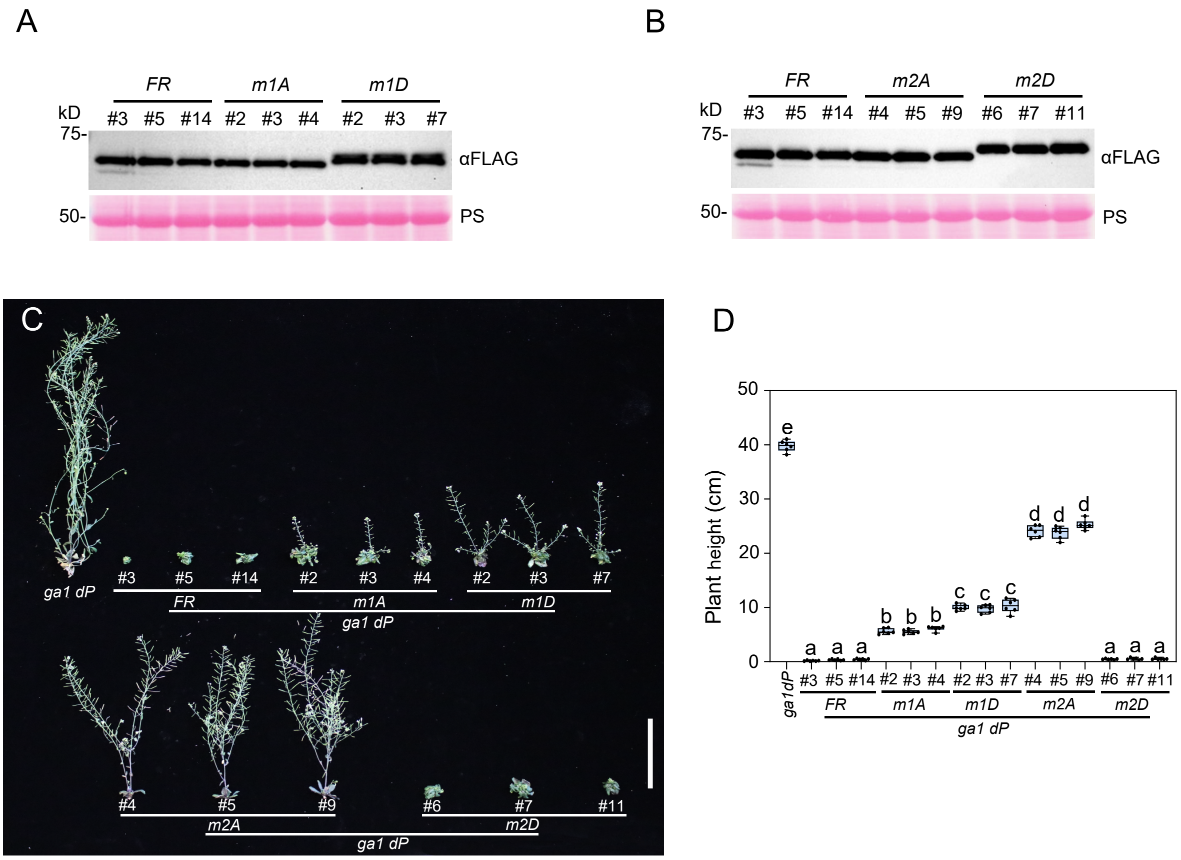


**Supplementary Figure 1.** Phenotype analysis of independent transgenic lines carrying $P_{RGA}$*:FLAG-RGA/rga*. Three independent homozygous lines for each transgene were included. **A-B)** All transgenic lines accumulated similar levels of FLAG-RGA/rga. Immunoblots containing total proteins from $P_{RGA}$*:FLAG-RGA/rga* lines (3 independent lines for each genotype) were probed with anti-FLAG antibody. Ponceau S (PS) staining showed similar loading amounts. **C)** Representative 55-d-old plants under LD conditions. Bar = 10 cm. **D)** Boxplot showing final heights of different lines as labeled. n=6. Center lines and box edges are medians and the lower/upper quartiles, respectively. Whiskers extend to the lowest and highest data points within 1.5× interquartile range (IQR) below and above the lower and upper quartiles, respectively. Different letters above the bars represent significant differences ($p < 0.01$) as determined by two-tailed Student's *t* tests. The phenotypic analysis was repeated two times with similar results.

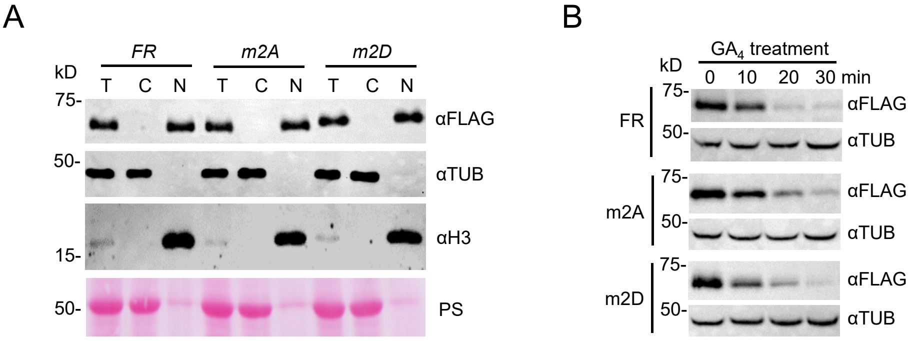


**Supplementary Figure 2.** rga$^{m2A}$ and rga$^{m2D}$ did not alter nuclear localization or GA-induced degradation of RGA in *Arabidopsis*. **A)** rga$^{m2A}$ and rga$^{m2D}$ did not alter nuclear localization. Immunoblot containing fractionated proteins from transgenic *Arabidopsis* (in *ga1 dP* background) carrying $P_{RGA}$:*FLAG-RGA/rga* was probed with anti-FLAG, anti-histone H3 and anti-tubulin (TUB) antibodies, separately. T, total protein; C, cytoplasmic fraction; N, nuclear fraction. **B)** rga$^{m2A}$ and rga$^{m2D}$ did not alter GA-induced degradation of RGA in *Arabidopsis*. Protein blots containing protein extracted from seedlings that were treated with 0.4 μM $GA_4$ at indicated time points were probed with anti-FLAG and anti-TUB antibodies (as the loading control), separately.

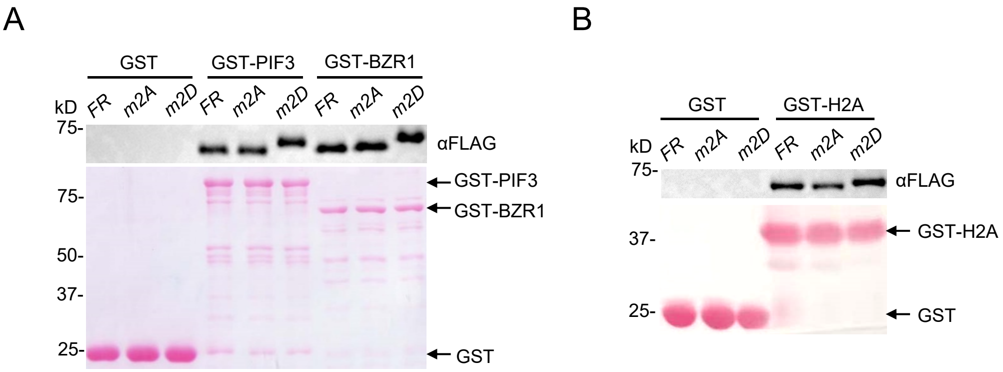


**Supplementary Figure 3.** In vitro pulldown assays to examine FLAG-RGA/rga interaction with TFs and H2A. **A-B)** The images of Ponceau S-stained blots show the GST and GST-BZR1, GST-PIF3, GST-H2A proteins used in the pull-down assays in Fig. 4A and 4B.

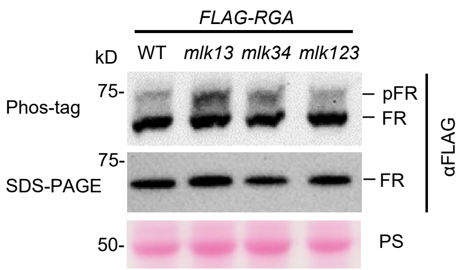


**Supplementary Figure 4.** RGA phosphorylation was not altered in *mlk* mutants. The protein blots contained total protein from $P_{RGA}$:*FLAG-RGA* in WT or *mlk* mutants as labeled. *mlk13*, homozygous *mlk1 mlk*3 double mutant; *mlk34*, homozygous *mlk3 mlk*4 double mutant; *mlk123*, homozygous *mlk1 mlk2 mlk*3 triple mutant. Proteins were analyzed by both standard SDS-PAGE and Phos-tag gels (containing 25 μM Phos-tag Acrylamide), followed by immunoblotting with an anti-FLAG antibody. pFR, phosphorylated FLAG-RGA. Ponceau S (PS)-stained blot indicated similar sample loading.
