## Supplementary Data for "Phosphorylation Promotes DELLA Activity by Enhancing Its Binding to Histone H2A at Target Chromatin in *Arabidopsis*"

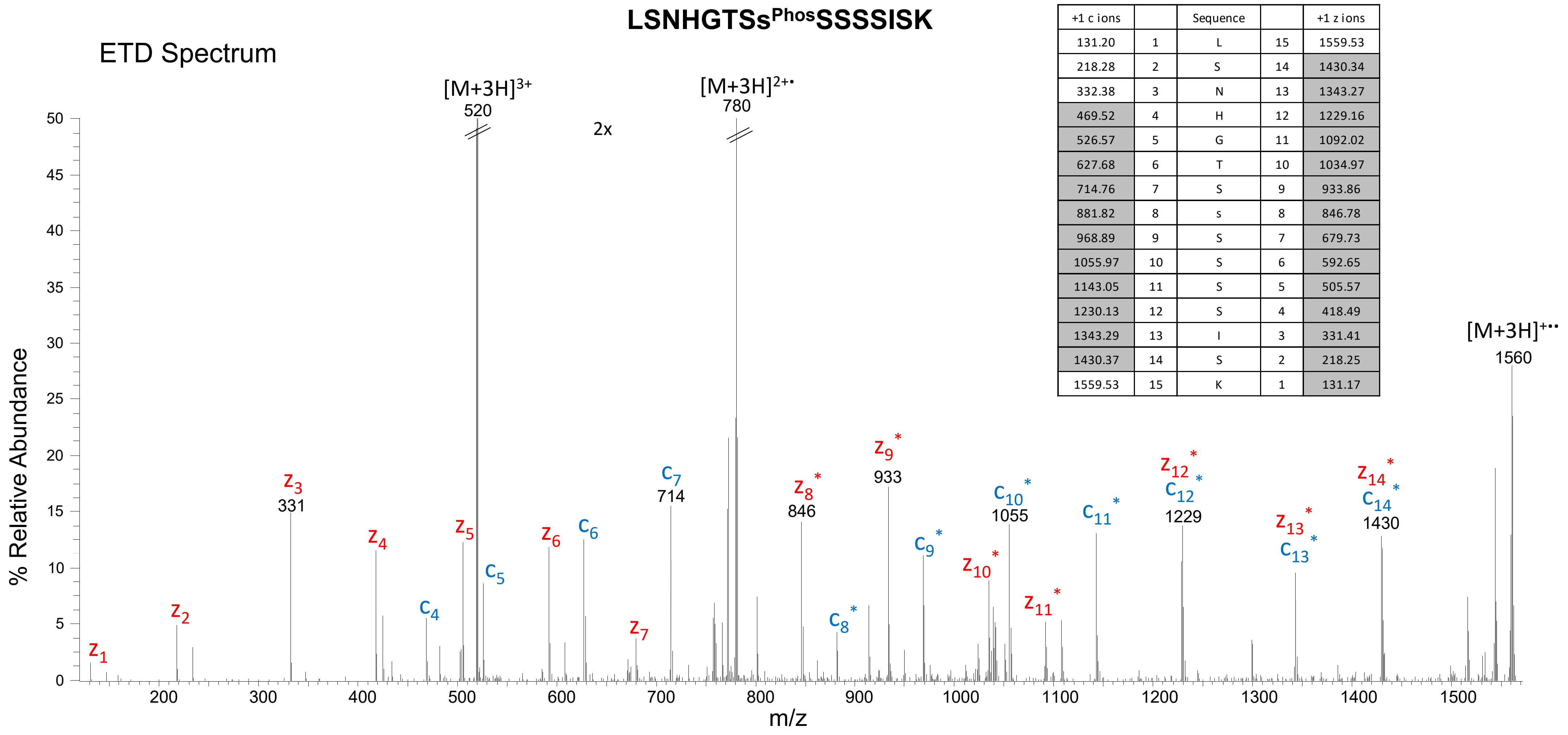


| +1 c ions | | Sequence | | +1 z ions |
|---|---|---|---|---|
| 131.20 | 1 | L | 15 | 1559.53 |
| 218.28 | 2 | S | 14 | 1430.34 |
| 332.38 | 3 | N | 13 | 1343.27 |
| 469.52 | 4 | H | 12 | 1229.16 |
| 526.57 | 5 | G | 11 | 1092.02 |
| 627.68 | 6 | T | 10 | 1034.97 |
| 714.76 | 7 | S | 9 | 933.86 |
| 881.82 | 8 | s | 8 | 846.78 |
| 968.89 | 9 | S | 7 | 679.73 |
| 1055.97 | 10 | S | 6 | 592.65 |
| 1143.05 | 11 | S | 5 | 505.57 |
| 1230.13 | 12 | S | 4 | 418.49 |
| 1343.29 | 13 | I | 3 | 331.41 |
| 1430.37 | 14 | S | 2 | 218.25 |
| 1559.53 | 15 | K | 1 | 131.17 |

**Supplementary Data 1. ETD MS2 spectrum of the RGA peptide LSNHGTSs$^{Phos}$SSSSISK (precursor ion m/z 520.23)** The spectrum displays average fragment ion masses detected with low resolution. Observed ions are highlighted in the fragment ion mass table. The data enable high-confidence identification of the peptide sequence and of the phosphorylation site, Ser8, as determined by the *-labeled fragment ions.

### LSNHGTSSs^Phos^SSSISKDK

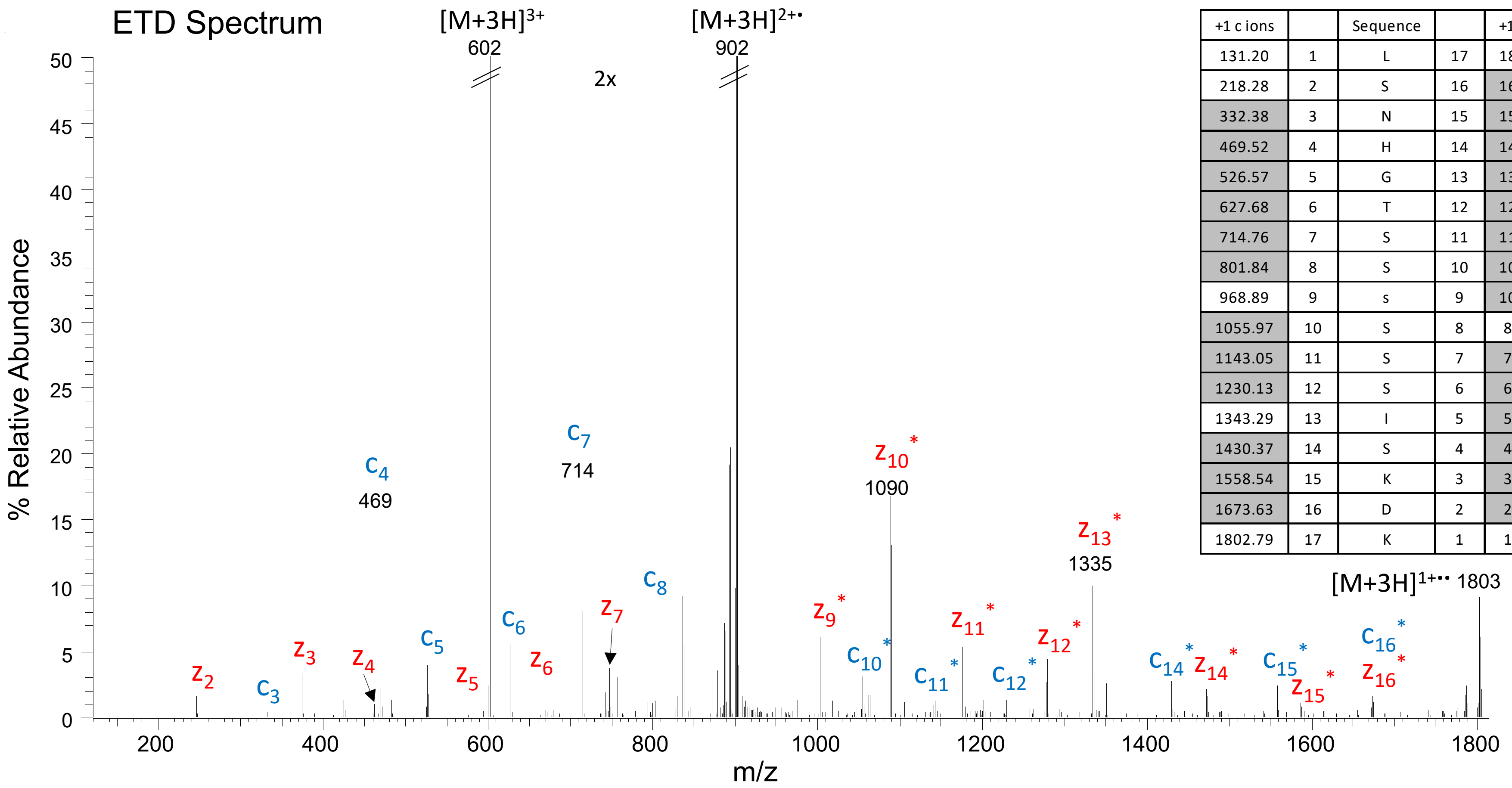


| +1 c ions | | Sequence | | +1 z ions |
|---|---|---|---|---|
| 131.20 | 1 | L | 17 | 1802.79 |
| 218.28 | 2 | S | 16 | 1673.61 |
| 332.38 | 3 | N | 15 | 1586.53 |
| 469.52 | 4 | H | 14 | 1472.42 |
| 526.57 | 5 | G | 13 | 1335.28 |
| 627.68 | 6 | T | 12 | 1278.23 |
| 714.76 | 7 | S | 11 | 1177.13 |
| 801.84 | 8 | S | 10 | 1090.05 |
| 968.89 | 9 | s | 9 | 1002.97 |
| 1055.97 | 10 | S | 8 | 835.91 |
| 1143.05 | 11 | S | 7 | 748.83 |
| 1230.13 | 12 | S | 6 | 661.75 |
| 1343.29 | 13 | I | 5 | 574.68 |
| 1430.37 | 14 | S | 4 | 461.52 |
| 1558.54 | 15 | K | 3 | 374.44 |
| 1673.63 | 16 | D | 2 | 246.26 |
| 1802.79 | 17 | K | 1 | 131.17 |

**Supplementary Data 2. ETD MS2 spectrum of the RGA peptide LSNHGTSSs^Phos^SSSISKDK (precursor ion m/z 601.27).** The spectrum displays average fragment ion masses detected with low resolution. Observed ions are highlighted in the fragment ion mass table. The data enable high-confidence identification of the peptide sequence and of the phosphorylation site, Ser9, as determined by the *-labeled fragment ions.

#### LSNHGTSSs$^{Phos}$SSs$^{Phos}$ISKDK

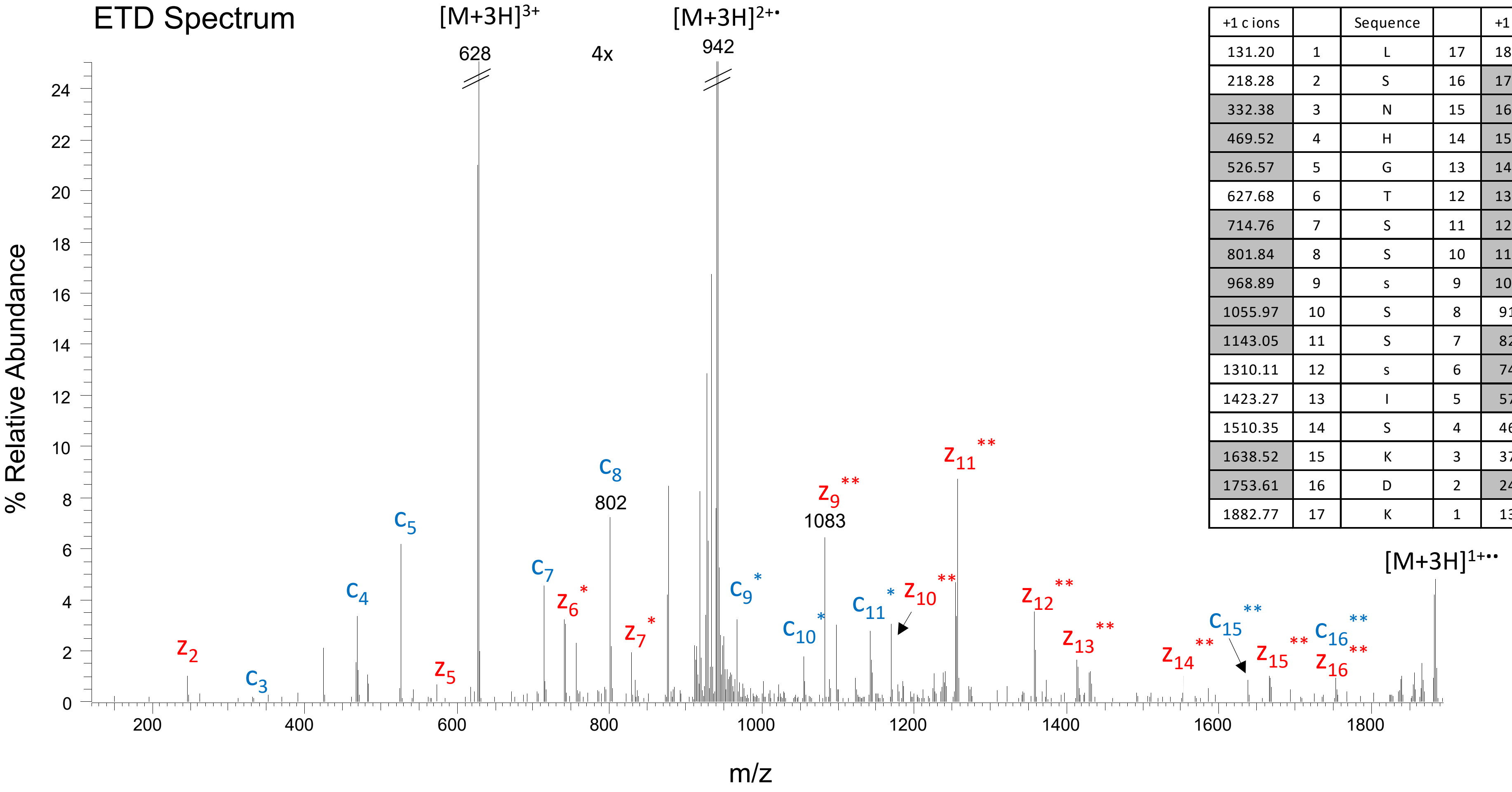


| +1 c ions | | Sequence | | +1 z ions |
|---|---|---|---|---|
| 131.20 | 1 | L | 17 | 1882.77 |
| 218.28 | 2 | S | 16 | 1753.59 |
| 332.38 | 3 | N | 15 | 1666.51 |
| 469.52 | 4 | H | 14 | 1552.40 |
| 526.57 | 5 | G | 13 | 1415.26 |
| 627.68 | 6 | T | 12 | 1358.21 |
| 714.76 | 7 | S | 11 | 1257.11 |
| 801.84 | 8 | S | 10 | 1170.03 |
| 968.89 | 9 | s | 9 | 1082.95 |
| 1055.97 | 10 | S | 8 | 915.89 |
| 1143.05 | 11 | S | 7 | 828.81 |
| 1310.11 | 12 | s | 6 | 741.73 |
| 1423.27 | 13 | I | 5 | 574.68 |
| 1510.35 | 14 | S | 4 | 461.52 |
| 1638.52 | 15 | K | 3 | 374.44 |
| 1753.61 | 16 | D | 2 | 246.26 |
| 1882.77 | 17 | K | 1 | 131.17 |

**Supplementary Data 3. ETD MS2 spectrum of the RGA peptide LSNHGTSSs$^{Phos}$SSs$^{Phos}$ISKDK (precursor ion m/z 627.93).** The spectrum displays average fragment ion masses detected with low resolution. Observed ions are highlighted in the fragment ion mass table. The data enable high-confidence identification of the peptide sequence and of the phosphorylation sites, Ser9 and Ser12, as determined by the *-labeled fragment ions.

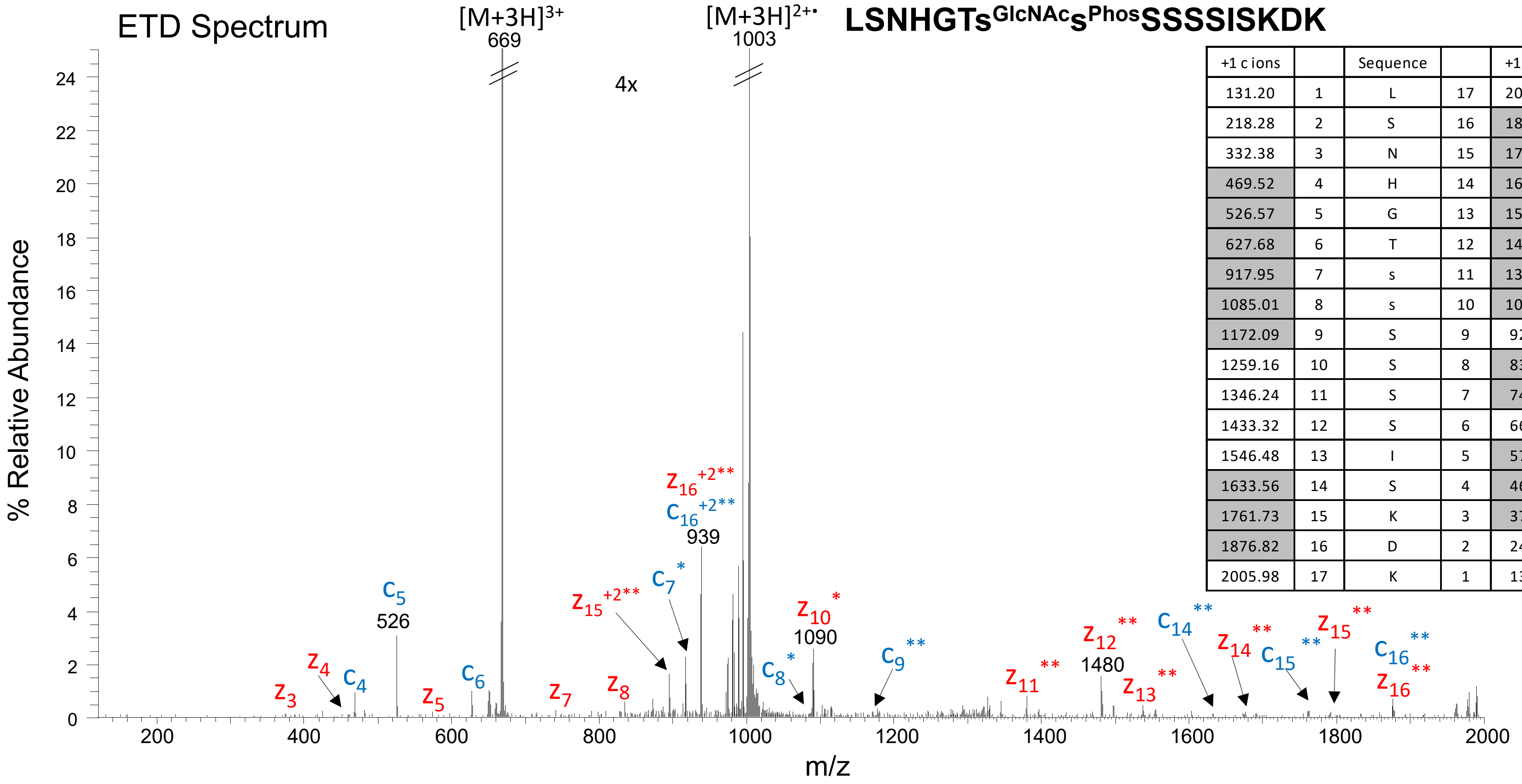


| +1 c ions | | Sequence | | +1 z ions |
|---|---|---|---|---|
| 131.20 | 1 | L | 17 | 2005.98 |
| 218.28 | 2 | S | 16 | 1876.80 |
| 332.38 | 3 | N | 15 | 1789.72 |
| 469.52 | 4 | H | 14 | 1675.62 |
| 526.57 | 5 | G | 13 | 1538.48 |
| 627.68 | 6 | T | 12 | 1481.42 |
| 917.95 | 7 | s | 11 | 1380.32 |
| 1085.01 | 8 | s | 10 | 1090.05 |
| 1172.09 | 9 | S | 9 | 922.99 |
| 1259.16 | 10 | S | 8 | 835.91 |
| 1346.24 | 11 | S | 7 | 748.83 |
| 1433.32 | 12 | S | 6 | 661.75 |
| 1546.48 | 13 | I | 5 | 574.68 |
| 1633.56 | 14 | S | 4 | 461.52 |
| 1761.73 | 15 | K | 3 | 374.44 |
| 1876.82 | 16 | D | 2 | 246.26 |
| 2005.98 | 17 | K | 1 | 131.17 |

**Supplementary Data 4. ETD MS2 spectrum of the RGA peptide LSNHGTs$^{GlcNAc}$s$^{Phos}$SSSSISKDK (precursor ion m/z 668.96).** The spectrum displays average fragment ion masses detected with low resolution. Observed ions are highlighted in the fragment ion mass table. The data enable high-confidence identification of the peptide sequence and of the O-GlcNAcylation site, Ser7, and phosphorylation site, Ser8, as determined by the *-labeled fragment ions.

**SCSs$^{Phos}$PDSMVTSTSTGTQIGK**

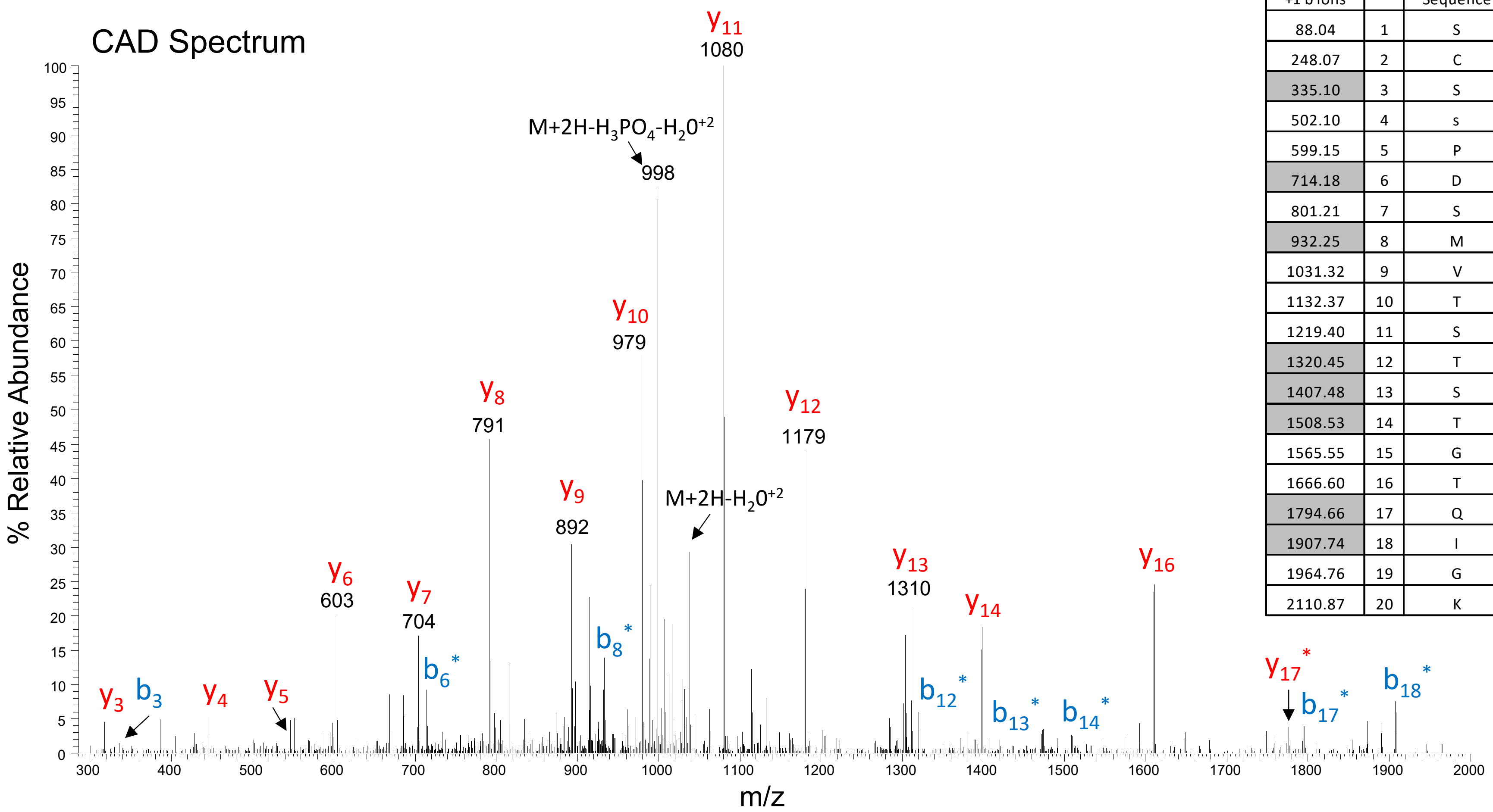


| +1 b ions | | Sequence | | +1 y ions |
|---|---|---|---|---|
| 88.04 | 1 | S | 20 | 2110.87 |
| 248.07 | 2 | C | 19 | 2023.83 |
| 335.10 | 3 | S | 18 | 1863.80 |
| 502.10 | 4 | s | 17 | 1776.77 |
| 599.15 | 5 | P | 16 | 1609.77 |
| 714.18 | 6 | D | 15 | 1512.72 |
| 801.21 | 7 | S | 14 | 1397.69 |
| 932.25 | 8 | M | 13 | 1310.66 |
| 1031.32 | 9 | V | 12 | 1179.62 |
| 1132.37 | 10 | T | 11 | 1080.55 |
| 1219.40 | 11 | S | 10 | 979.51 |
| 1320.45 | 12 | T | 9 | 892.47 |
| 1407.48 | 13 | S | 8 | 791.42 |
| 1508.53 | 14 | T | 7 | 704.39 |
| 1565.55 | 15 | G | 6 | 603.35 |
| 1666.60 | 16 | T | 5 | 546.32 |
| 1794.66 | 17 | Q | 4 | 445.28 |
| 1907.74 | 18 | I | 3 | 317.22 |
| 1964.76 | 19 | G | 2 | 204.13 |
| 2110.87 | 20 | K | 1 | 147.11 |

**Supplementary Data 5. CAD MS2 spectrum of the RGA peptide SCSs$^{Phos}$PDSMVTSTSTGTQIGK (precursor ion m/z 1055.94).** The spectrum displays average fragment ion masses detected with low resolution. Observed ions are highlighted in the fragment ion mass table. The data enable high-confidence identification of the peptide sequence and of the phosphorylation site, Ser4, as determined by the *-labeled fragment ions.
