## Supplementary Table 1 for "Phosphorylation Promotes DELLA Activity by Enhancing Its Binding to Histone H2A at Target Chromatin in *Arabidopsis*"

**Supplementary Table 1. Posttranslational modifications in FLAG-RGA$^{GKG}$**

| Residues[a] | Peptide sequence[b] | PTM | Sites[c] | Relative Abundance[d] | | | | |
|---|---|---|---|---|---|---|---|---|
| | | | | *ga1-3* | *sly1-10* | *sly1-10 spy-12* | *sly1-10 spy-19* | *sly1-10 sec-3* |
| 12-26 | LSNHGt$^{GlcNAc}$SSSSSSISK | 1 GlcNAc | T17 | 1.6% | 2.1% | 8.1% | 10.7% | 0.0% |
| 12-26(28) | LSNHGTs$^{GlcNAc}$SSSSSISK(DK) | 1 GlcNAc | S18 | | | | | |
| 12-26(28) | LSNHGTSs$^{GlcNAc}$SSSSISK(DK) | 1 GlcNAc | S19 | | | | | |
| 12-28 | LSNHGTS[SSSSS]$^{GlcNAc}$ISKDK | 1 GlcNAc | S[19-23] | | | | | |
| 12-26(28) | LSNHGTs$^{GlcNAc}$s$^{Phos}$SSSSISKDK | 1 GlcNAc + 1 Phos | S18, S19 | 0.3% | 0.0% | 0.0% | 0.0% | 0.0% |
| 12-28 | LSNHGt$^{Fuc}$SSSSSSISKDK | 1 Fuc | T17 | 0.3% | 0.7% | 0.0% | 0.0% | 2.0% |
| 12-28 | LSNHGTSSs$^{Fuc}$SSSISKDK | 1 Fuc | S20 | | | | | |
| 12-28 | LSNHGT[SS]$^{Fuc}$SSSSISKDK | 1 Fuc | S[18-19] | | | | | |
| 12-26 | LSNHGTs$^{Hex}$SSSSSISK | 1 Hex | S18 | 0.0% | 0.1% | 0.0% | 0.1% | 0.0% |
| 12-26 | LSNHG[TSSS]$^{Hex}$SSSISK | 1 Hex | [T17-S20] | | | | | |
| 12-26(28) | LSNHGTSs$^{Phos}$SSSSISK(DK) | 1 Phos | S19 | 8.9% | 4.1% | 2.2% | 3.2% | 7.8% |
| 12-28 | LSNHGTSSs$^{Phos}$SSSISKDK | 1 Phos | S20 | | | | | |
| 12-28 | LSNHGT[SSSSSS]$^{Phos}$ISKDK | 1 Phos | S[18-23] | | | | | |
| 12-26(28) | L[SNHGTSSSSSS]$^{Phos}$ISK(DK) | 1 Phos | S13 or [T17-S23] | | | | | |
| 12-26 | LSNHGTSs$^{Phos}$SSSs$^{Phos}$ISK | 2 Phos | S19, S23 | 17.4% | 1.1% | 0.2% | 0.3% | 4.0% |
| 12-28 | LSNHGTSSs$^{Phos}$SSs$^{Phos}$ISKDK | 2 Phos | S20, S23 | | | | | |
| 12-28 | LSNHGTS[SSSSS]$^{2\ Phos}$ISKDK | 2 Phos | S[19-23] | | | | | |
| 12-28 | L[SNHGTSSSSSS]$^{2\ Phos}$ISKDK | 2 Phos | Not mapped to specific residue | | | | | |
| 12-28 | L[SNHGTSSSSSS]$^{3\ Phos}$ISKDK | 3 Phos | Not mapped to specific residue | 4.2% | 0.1% | 0.0% | 0.0% | 0.5% |
| 140-164 | VIPGNAIYQFPAIDS[SSSS]$^{Phos}$NNQNKR | 1 Phos | S[155-158] | 2.0% | 0.0% | 0.1% | 0.4% | 0.3% |
| 140-164 | VIPGNAIYQFPAID[SSSSS]$^{2\ Phos}$NNQNKR | 2 Phos | S[154-158] | 0.4% | 0.0% | 1.0% | 0.0% | 0.0% |
| 165-185 | LKSC[SSPDS]$^{GlcNAc}$MVTSTSTGTQIGK | 1 GlcNAc | S[169-170] or S173 | 3.7% | 4.2% | 4.1% | 5.4% | 0.0% |
| 165-185 | LK[SCSSPDSMVTSTSTGT]$^{GlcNAc}$QIGK | 1 GlcNAc | Not mapped to specific residue | | | | | |
| 167-185 | SCSSPDSMVt$^{GlcNAc}$STSTGTQIGK | 1 GlcNAc | T176 | | | | | |
| 167-185 | [SCSSPDSMVTSTSTGT]$^{GlcNAc}$QIGK | 1 GlcNAc | Not mapped to specific residue | | | | | |
| 165-185 | LK[SCSSPDSMVTSTSTGT]$^{Hex}$QIGK | 1 Hex | Not mapped to specific residue | 0.1% | 0.7% | 0.1% | 0.2% | 0.0% |
| (165)167-185 | (LK)SCSs$^{Phos}$PDSMVTSTSTGTQIGK | 1 Phos | S170 | 27.2% | 23.4% | 15.6% | 21.8% | 24.0% |
| 165-185 | LK[SCSSPDS]$^{Phos}$MVTSTs$^{Phos}$TGTQIGK | 2 Phos | S[167, 169, 170 or 173], S179 | 1.4% | 0.0% | 0.0% | 0.0% | 0.0% |
| 165-185 | LK[SCSSPDS]$^{2\ Phos}$MVTSTSTGTQIGK | 2 Phos | S[167, 169, 170 or 173] | | | | | |
| 186-207 | GVIGTTV[TTTTTT]$^{GlcNAc}$TTAAGESTR | 1 GlcNAc | Not mapped to specific residue | 44.1% | 46.4% | 49.7% | 44.1% | 0.1% |
| 186-207 | GVIGTTV[TTTT]$^{2\ GlcNAc}$TTTTAAGESTR | 2 GlcNAc | Not mapped to specific residue | 13.5% | 7.4% | 7.4% | 12.2% | 0.0% |
| 186-207 | GVIG[TTVTTT]$^{2\ GlcNAc}$TTTTTAAGESTR | 2 GlcNAc | Not mapped to specific residue | | | | | |
| 186-207 | GVIG[TTVTTTTTTTTAAGEST]$^{3\ GlcNAc}$R | 3 GlcNAc | Not mapped to specific residue | 3.4% | 0.6% | 0.8% | 2.1% | 0.0% |
| 186-207 | GVIG[TTVTTTTTTTTAAGEST]$^{4\ GlcNAc}$R | 4 GlcNAc | Not mapped to specific residue | 0.5% | 0.0% | 0.0% | 0.0% | 0.0% |
| 186-207 | GVIG[TTVTTTTTTTTAAGEST]$^{Fuc}$R | 1 Fuc | Not mapped to specific residue | 0.0% | 0.1% | 0.0% | 0.0% | 0.3% |
| 186-207 | GVIGTTV[TT]$^{Hex}$TTTTTTAAGESTR | 1 Hex | Not mapped to specific residue | 6.6% | 8.6% | 7.2% | 5.7% | 0.0% |
| 186-207 | GVIG[TTVTTTTTTTTAAGEST]$^{2\ Hex}$R | 2 Hex | Not mapped to specific residue | 0.3% | 0.3% | 0.5% | 0.6% | 0.0% |
| 186-207 | GVIG[TTVTTTTTTTTAAGEST]$^{GlcNAc\ +\ Fuc}$R | 1 GlcNAc + 1 Fuc | Not mapped to specific residue | 0.4% | 0.2% | 0.0% | 0.0% | 0.0% |
| 186-207 | GVIG[TTVTTTTTTTT]$^{GlcNAc\ +\ Hex}$AAGESTR | 1 GlcNAc + 1 Hex | Not mapped to specific residue | 5.6% | 3.4% | 4.2% | 6.4% | 0.0% |
| 186-207 | GVIG[TTVTTTTTTTT]$^{2\ GlcNAc\ +\ Hex}$AAGESTR | 2 GlcNAc + 1 Hex | Not mapped to specific residue | 1.8% | 0.4% | 0.5% | 1.6% | 0.0% |
| 186-207 | GVIG[TTVTTTTTTTT]$^{GlcNAc\ +\ 2\ Hex}$AAGESTR | 1 GlcNAc + 2 Hex | Not mapped to specific residue | 0.3% | 0.0% | 0.0% | 0.1% | 0.0% |

[a] Residues in WT RGA, and numbers in parentheses indicate the residues of longer peptides resulting from incomplete digestion, and abundances of the short and long peptides were combined. [b] Underlined "K" refers to the engineered Lys in FLAG-RGA$^{GKG}$; when a specific amino acid could not be mapped, brackets indicate the region in the peptide that contains the PTM(s). [c] Site was determined by correct elution profile, accurate mass, and signature CAD and/or ETD fragmentation. [d] Peptide abundances were determined from ion currents taken from the MS1 survey scan. PTM levels are reported as the % of the total peptide abundance detected. Total abundances were calculated from ion currents observed in the MS1 mass spectra [(modified peptide ion current)/ (sum of all modified peptides + unmodified peptide ion current)] x 100. A 0.0% relative abundance corresponds to less than 0.1% relative abundance. The percentage values are average of three biological repeats for *ga1-3* and *sly1-10* or two biological repeats for *sly1-10 sec-3*.
