## Supplementary Table 2 for "Phosphorylation Promotes DELLA Activity by Enhancing Its Binding to Histone H2A at Target Chromatin in *Arabidopsis*"

**Supplementary Table 2. List of Primers**

| Name | Sequence | Use | Note |
|---|---|---|---|
| SCL3-F | ACACCCAAGCCTCAGCCTCATCTC | ChIP-qPCR | AT1G50420, Ref.1 |
| SCL3-R | GGAATCATGACTATATATTTCTACATC | | |
| GID1B-F | GAGAGGGTCCCTGACT | ChIP-qPCR | AT3G63010, Ref.1 |
| GID1B-R | AGAGCAAGGAAGGAACG | | |
| PP2A-F | CGGCTTTCATGATTCCCTCT | ChIP-qPCR (reference gene) | At1g13320, Ref.3 |
| PP2A-R | GCCTTAAGCTCCGTTTCCTACTT | | |
| IAA16-F | CATTGATCGGAAGGAATGTTT | ChIP-qPCR | AT3G04730, Ref.1 |
| IAA16-R | TGCCCCTGTGGCCTTGTCTTGG | | |
| EXP8-F | GTTGATCACGTTTAGGCACTTA | ChIP-qPCR | AT2G40610, Ref.1 |
| EXP8-R | AGACAGGCCACTAATACAT | | |
| SCL3-qF | CAGCTGAGGCACGTGAGAATGAT | RT-qPCR | AT1G50420, Ref.2 |
| SCL3-qR | ACCACCATGACCTTTGGAGACAAAC | | |
| GID1B-qF | AGAGGTCAAAGCCTTAAAGGAGTC | RT-qPCR | AT3G63010, Ref.2 |
| GID1B-qR | CTTCAAGACCAGTCTTCTTAAGCCC | | |
| IAA16-qF | TGGATGCTTGTAGGAGAC | RT-qPCR | AT3G04730 |
| IAA16-qR | AGTCCGATTGCTTCTGAT | | |
| EXP8-qF | CATGTATGAAGAAAGGAGGAATAAG | RT-qPCR | AT2G40610, Ref.2 |
| EXP8-qR | AACTGCCAATTAGAAGGAGCCACG | | |
| PP2A-qF | TATCGGATGACGATTCTTCGTGCAG | RT-qPCR (reference gene) | At1g13320, Ref. 3 |
| PP2A-qR | GCTTGGTCGACTATCGGAATGAGAG | | |
| RGA-prom-400-F | CCTTATCACTTTAGATGTGTATG | For overlap PCR mutagenesis of 6xHis-3xFLAG-RGA | |
| RGA-Mut2-R | AGCCGCAGCCCCGTGGTTGGACAATCGACC | For overlap PCR mutagenesis of 6xHis-3xFLAG-RGA | rga-m1A |
| RGA-Mut7-F | CACGGGGCTGCGGCTGCAGCTGCGGCAATCGCTAAAGATAAGATGATGATG | For overlap PCR mutagenesis of 6xHis-3xFLAG-RGA | |
| RGA-Mut3-R | ATCATCTTCCCCGTGGTTGGACAATCGACC | For overlap PCR mutagenesis of 6xHis-3xFLAG-RGA | rga-m1D |
| RGA-Mut8-F | CACGGGGAAGATGATGATGACGATGACATCGATAAAGATAAGATGATGATG | For overlap PCR mutagenesis of 6xHis-3xFLAG-RGA | |
| RGA-minusB2-R | CTCAGTACGCCGCCGTCGAGAGTTTC | For overlap PCR mutagenesis of 6xHis-3xFLAG-RGA | |
| RGA-3xFLAG-B1F-1 | GGGGACAAGTTTGTACAAAAAAGCAGGCTCTCATAGCGGTACCTGAAAAATGAGA | Cloning of 6xHis-3xFLAG-RGA into pDONR207 | |
| RGA-B2R-1 | GGGGACCACTTTGTACAAGAAAGCTGGGTCTCAGTACGCCGCCGTCGAGAGTTTC | Cloning of 6xHis-3xFLAG-RGA into pDONR207 | |
| rgam2A-F | GCTGCTCCTGATGCTATGGTTGCTGCTGCTGCTGCTGGTACGCAGATTGGTGGAGTC | For overlap PCR mutagenesis of 6xHis-3xFLAG-RGA | rga-m2A |
| rgam2A-R | AGCAGCAGCAGCAGCAACCATAGCATCAGGAGCAGCGCATGATTTCAAACGCTTGTTCTG | | |
| rgam2D-F | GACGATCCTGATGATATGGTTGAAGACGAAGATGAGGGTACGCAGATTGGTGGAGTC | For overlap PCR mutagenesis of 6xHis-3xFLAG-RGA | rga-m2D |
| rgam2D-R | CTCATCTTCGTCTTCAACCATATCATCAGGATCGTCGCATGATTTCAAACGCTTGTTCTG | | |
