## Supplementary Table 3 for "Phosphorylation Promotes DELLA Activity by Enhancing Its Binding to Histone H2A at Target Chromatin in *Arabidopsis*"

**Supplementary Table 3. List of Plasmids**

| Plasmid | Source vector (Reference) | Source insert (Reference) | Cloning method | Note |
|---|---|---|---|---|
| GST | pGEX-4T-1 (GE) | | | *E. coli.* GST |
| GST-PIF3 | Ref.1 | PIF3 | | *E. coli.* GST-PIF3 |
| GST-BZR1 | pENTR1A-HA-BZR1, Ref. 1 | BZR1 | | *E. coli.* GST-BZR1 |
| GST-H2A | pGEXM, Ref.2 | H2A | | *E. coli.* GST-H2A |
| pDONR207-6xHis-3xFLAG-rga-m1A | pDONR207 (Thermo-Fisher) | Overlapping PCR, RGA-prom-400-F/RGA-Mut2-R and RGA-Mut7-F/RGA-minusB2-R; then RGA-3xFLAG-B1F-1/RGA-B2R-1 | Overlapping PCR and Gateway cloning BP reaction | Entry vector |
| pDONR207-6xHis-3xFLAG-rga-m1D | pDONR207 (Thermo-Fisher) | Overlapping PCR, RGA-prom-400-F+RGA-Mut3-R and RGA-Mut8-F+RGA-minusB2-R; then RGA-3xFLAG-B1F-1/RGA-B2R-1 | Overlapping PCR and Gateway cloning BP reaction | Entry vector |
| pDONR207-6xHis-3xFLAG-rga-m2A | pDONR207 (Thermo-Fisher) | Overlapping PCR, RGA-prom-400-F+rgam2A-R and rgam2A-F+RGA-minusB2-R; then RGA-3xFLAG-B1F-1/RGA-B2R-1 | Overlapping PCR and Gateway cloning BP reaction | Entry vector |
| pDONR207-6xHis-3xFLAG-rga-m2D | pDONR207 (Thermo-Fisher) | Overlapping PCR, RGA-prom-400-F+rgam2D-R and rgam2D-F+RGA-minusB2-R; then RGA-3xFLAG-B1F-1/RGA-B2R-1 | Overlapping PCR and Gateway cloning BP reaction | Entry vector |
| pBm43GW | Ref. 3 | | | Multi-Gateway cloning destination vector |
| JO35 | pDONR P4-P1R (Thermo-Fisher) | RGA promoter,8.1kb, gateway cloning for RGA promoter | Gateway cloning BP reaction | gift from Neil Olszewski |
| JO15 | pDONR P2R-P3 (Thermo-Fisher) | RGA 3'UTR, gateway cloning for RGA 3'UTR | | gift from Neil Olszewski |
| pRGA-6xHis-3xFLAG-rga-m1A | pBm43GW. Ref. 3 | JO35/pDONR207-6xHis-3xFLAG-rga-m1A/JO15 | Multi-Gateway cloning LR reaction | $P_{RGA}$:*FLAG-rga*$^{m1A}$ for Arabidopsis transformation |
| pRGA-6xHis-3xFLAG-rga-m1D | pBm43GW. Ref. 3 | JO35/pDONR207-6xHis-3xFLAG-rga-m1D/JO15 | Multi-Gateway cloning LR reaction | $P_{RGA}$:*FLAG-rga*$^{m1D}$ for Arabidopsis transformation |
| pRGA-6xHis-3xFLAG-rga-m2A | pBm43GW. Ref. 3 | JO35/pDONR207-6xHis-3xFLAG-rga-m2A/JO15 | Multi-Gateway cloning LR reaction | $P_{RGA}$:*FLAG-rga*$^{m2A}$ for Arabidopsis transformation |
| pRGA-6xHis-3xFLAG-rga-m2D | pBm43GW. Ref. 3 | JO35/pDONR207-6xHis-3xFLAG-rga-m2D/JO15 | Multi-Gateway cloning LR reaction | $P_{RGA}$:*FLAG-rga*$^{m2D}$ for Arabidopsis transformation |
